# Species-specific assembly of root-associated bacterial microbiota mediated by a combination of plant specialized metabolites

**DOI:** 10.1101/2020.09.25.313031

**Authors:** Tomohisa Shimasaki, Sachiko Masuda, Ruben Garrido-Oter, Takashi Kawasaki, Yuichi Aoki, Arisa Shibata, Wataru Suda, Ken Shirasu, Kazufumi Yazaki, Ryohei Thomas Nakano, Akifumi Sugiyama

## Abstract

Plant specialized metabolites (PSMs) influence the taxonomic compositions of the root-associated microbiota; however, the underlying molecular mechanisms and evolutionary trajectories remain elusive. Here, we show that the bacterial genus *Arthrobacter* is predominant in the tobacco endosphere, and that its enrichment is mediated by a combination of two tobacco-specific PSMs, santhopine and nicotine The isolation and whole genome sequencing of a representative set of *Arthrobacter* strains identified independent genomic features, including but not limited to genes for santhopine and nicotine catabolism, which are associated with the colonization competence of tobacco roots. Taken together, these data suggest that plant species-specific root microbiota assembly is mediated by bacterial catabolism of a cocktail of PSMs synthesized by the host plant.

## Introduction

Plant roots secrete their photosynthates to the rhizosphere, a fraction of soil surrounding the root (*1*), creating a nutrient-rich environment with a distinctive metabolic profile compared to the bulk soil (*2, 3*). These rhizosphere metabolites attract or repel particular soil bacteria, resulting in characteristic bacterial communities in the rhizosphere. A subset of rhizosphere bacteria can further colonize the root surface and the interior, constituting the rhizoplane and endosphere bacterial communities, respectively, which are collectively called the root microbiota (*4*). Four bacterial phyla, Proteobacteria, Actinobacteria, Bacteroidetes, and Firmicutes, dominate the root microbiota (*5–7*), whereas each plant species harbors a different composition of root microbiota at lower taxonomic levels (*8*), suggesting the ability of plants to modulate root microbiota in a manner specific to each plant lineage.

Plant specialized metabolites (PSMs) play important roles in the interaction between the host and its root microbiota, affecting its taxonomic composition (*9*). For instance, the disruption of the triterpene or coumarin biosynthetic pathway in *Arabidopsis thaliana* resulted in different taxonomic structures of the root microbiota (*10–12*). Soil treatment with isoflavone or soyasaponin, two major PSMs in soybean root exudates, altered the compositions of the soil bacterial community and enriched bacteria commonly found in soybean roots (*13, 14*). Recent large-scale comparative genomics analyses also showed that the bacterial species inhabiting plant tissues had evolved characteristic functions, including specific metabolic capacities, which enabled their adaptation to plants (*15*). However, the mechanisms by which PSMs affect the relative abundance of a given bacterial species and whether single or multiple classes of PSMs are needed to shape the microbial community remains unclear. The evolutionary processes through which plants and bacteria establish such PSM-mediated interactions also remain to be addressed.

We employed *Nicotiana tabacum* (cultivated tobacco), whose roots accumulate characteristic PSMs, such as santhopine and nicotine (Fig. 1A) (*16, 17*). Santhopine is an opine, whose biosynthesis is induced in crown gall tumors and hairy roots upon infection by pathogenic *Rhizobium* strains (*18, 19*). During the diversification of the *Nicotiana* genus, some of its species acquired genes for opine biosynthesis, including the *mannopine synthase 2* (*mas2*) gene, which encodes the enzyme for santhopine synthesis, likely *via* a horizontal gene transfer (HGT) event from *Rhizobium* species (*20–22*). *N. tabacum* then inherited the *mas2* gene from its parental wild species *Nicotiana tomentosiformis* (*23*). Opine-catabolic genes have been found in a limited member of bacterial species (*24*), suggesting that the opines synthesized by plants may serve as nutrients for specific groups of bacteria. In contrast, nicotine, a major alkaloid of *Nicotiana* species, exhibits strong toxicity and contributes to chemical defense against insect predators (*25–27*). Both santhopine and nicotine are catabolized by several bacterial species (*28, 29*), implying the possible involvement of those metabolites in the interaction between tobacco and its root-associated microbiota.

**Fig. 1.**
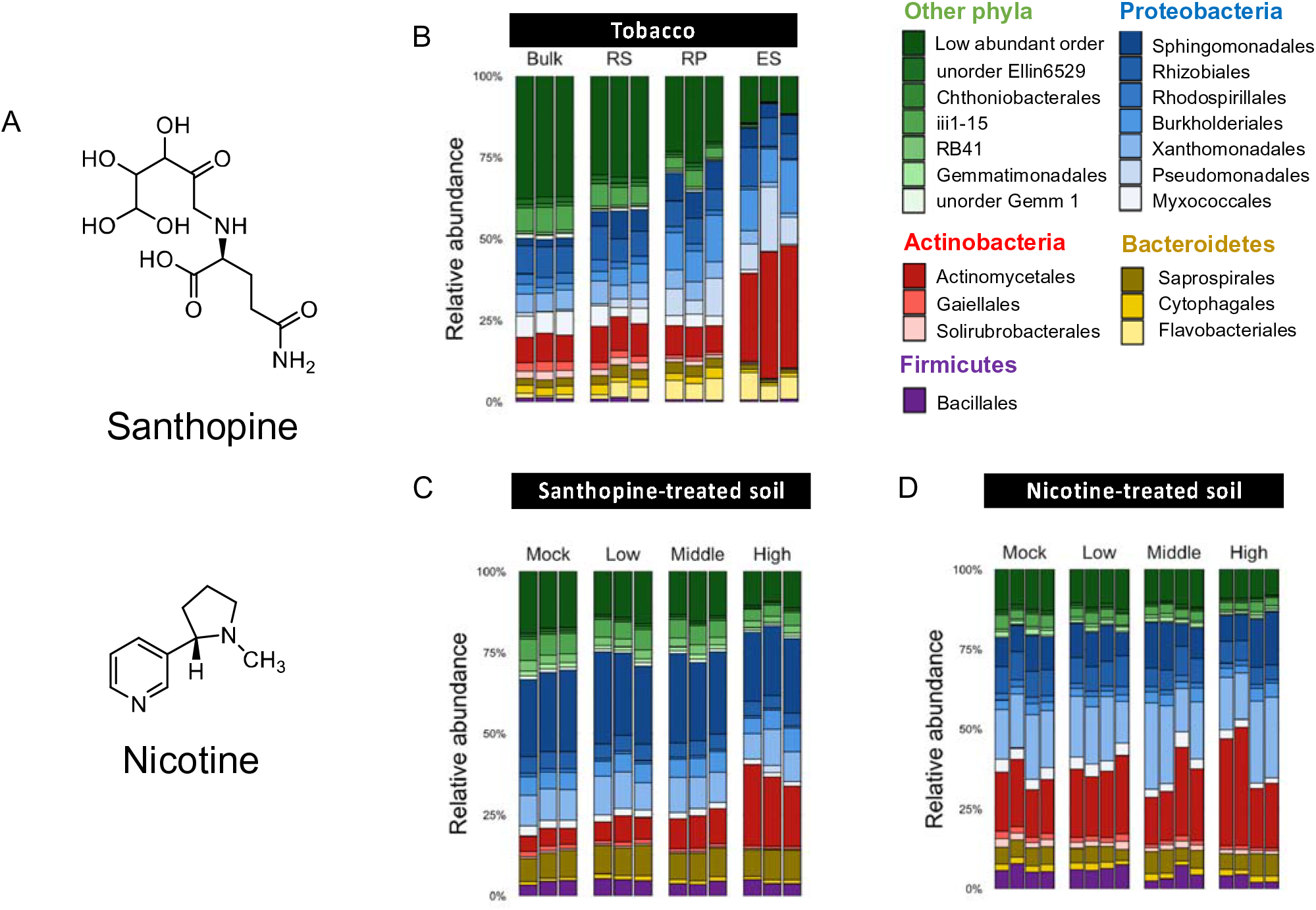
Bacterial community composition in tobacco roots and santhopine-/nicotine-treated soil. **(A)** Chemical structure of santhopine and nicotine. **(B-D)** Composition of the 20 most abundant bacterial orders across samples in tobacco roots **(B)**, santhopine-treated soil **(C)**, and nicotine-treated soil **(D)**. Bulk, bulk soil; RS, rhizosphere; RP, rhizoplane; ES, endosphere; Mock, bulk soils treated with sterile water; Low, middle, and high, bulk soils treated with 50, 250, and 1000 nmol g^−1^ soil of santhopine or nicotine, respectively.

In this study, we analyzed the bacterial communities of the tobacco endosphere and the effect of santhopine and nicotine on the soil bacterial community. This led us to focus on the genus *Arthrobacter*, which was specifically enriched in the tobacco endosphere and PSM-treated soils. By integrating culture-dependent characterization and comparative genomics of *Arthrobacter* isolates, we herein propose a model that explains a process of host-specific root microbiota assembly mediated by bacterial catabolism of a combination of PSMs synthesized by the host plant.

## Results

### *Arthrobacter* is predominantly found in the tobacco endosphere and enriched in soils by santhopine and nicotine treatment

We investigated the taxonomic composition of the bacterial community associated with tobacco roots by sequencing the V4 region of bacterial *16S* rRNA genes. Analyses of within-sample diversity [Shannon diversity index and the number of observed amplicon sequence variants (ASVs)] revealed that the α-diversity of the bacterial population gradually decreased from bulk soil to endosphere (Fig. S1A and B). Analysis of taxonomic profiles across samples revealed the distinctive community structure of the endosphere compared to rhizosphere and rhizoplane fractions (*P* = 1.00 × 10^−3^; Fig. 1B; Fig. S2). Bacterial members belonging to the order Actinomycetales were significantly enriched in the endosphere compared to the rhizosphere and rhizoplane fractions (*P* = 6.39 × 10^−12^ and 1.59 × 10^−10^, respectively). In parallel, to specifically quantify the effects of santhopine and nicotine, we treated the same soils with either of these metabolites at three different concentrations and compared the compositions of the bacterial community (Fig. 1C and D). Santhopine treatment enriched Actinomycetales in a dose-dependent manner (*P* = 1.78 × 10^−3^ for the highest concentration compared to the mock control; Fig. 1C), whereas nicotine treatment only moderately shifted the overall taxonomic composition (*P* = 6.10 × 10^−2^ for the highest concentration compared to the mock control; Fig. 1D). The Shannon diversity index of both santhopine- and nicotine-treated soils decreased along with the chemical concentrations of these compounds (Fig. S1C and E), and the number of observed ASVs was lower in santhopine-treated soil but not in nicotine-treated soils (Fig. S1D and F). Together, these data suggest that santhopine and nicotine have significant but different effects on soil bacterial community composition.

Interestingly, the bacterial community composition of mock-treated soils was more similar to bulk soils than the tobacco endosphere based on weighted UniFrac distance (Fig. S3A and C). In contrast, both santhopine and nicotine treatment shifted the soil bacterial community profiles toward the tobacco endosphere (Fig. S3B and D). Fourteen bacterial families, mainly belonging to the phyla Proteobacteria and Actinobacteria, were enriched in the tobacco endophytic compartment compared to the rhizoplane, and three and five bacterial families were enriched by santhopine and nicotine treatment, respectively (Fig. 2A). Among these families, Micrococcaceae was the only one commonly enriched in the tobacco endosphere, santhopine-treated soil, and nicotine-treated soil. At the genus level, all sequence reads classified as belonging to the family Micrococcaceae were assigned to the genus *Arthrobacter,* which represented about 15% of the total bacterial sequences in both the tobacco endosphere and santhopine- and nicotine-treated soils (Fig. 2B). Enrichment of *Arthrobacter* in the endosphere is unique to tobacco plants and was not observed in tomato (S*olanum lycopersicum*), soybean (*Glycine max*), or bitter melon (*Momordica charantia*) plants grown in the same soil. Given that the production of santhopine and nicotine is specific to tobacco plants, these results suggest a crucial role of these metabolites in forming tobacco-specific root microbiota structures.

**Fig. 2.**
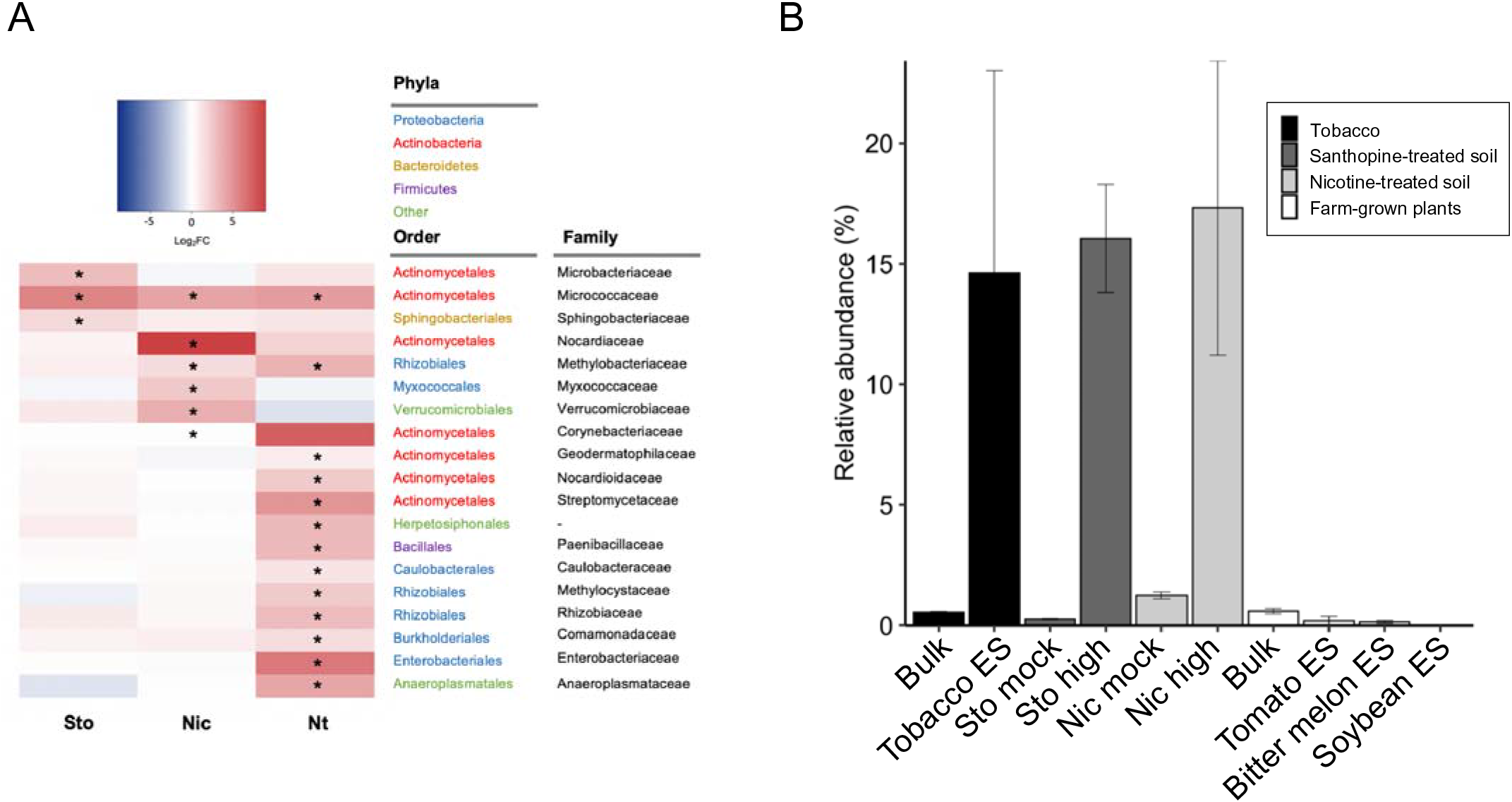
Differential abundance analysis of each sample. **(A)** Heatmap showing the fold changes at the log_2_ scale in the tobacco endosphere and santhopine- and nicotine-treated soils at the highest concentrations, compared to rhizoplane, santhopine mock, and nicotine mock treatments, respectively. Asterisks indicate statistical significance (false discovery rate < 0.05). Nic, nicotine-treated soil; Sot, santhopine-treated soil; Nt, tobacco endosphere. **(B)** Mean relative abundance of the genus *Arthrobacter* in soils treated with santhopine or nicotine and in the tobacco, tomato, bitter melon, and soybean root endosphere along with their respective bulk soil samples. Error bars represent standard deviation (n = 3 for tobacco roots and santhopine-treated soil and n = 4 for nicotine-treated soil).

### Predominance of a monophyletic clade of the genus *Arthrobacter* in the tobacco endosphere

To investigate the taxonomic diversity within the genus *Arthrobacter*, we isolated 252 individual bacterial strains from surface-sterilized tobacco roots, which corresponded to the endosphere fraction in the community profiling experiments, as well as from santhopine or nicotine-treated soil. These isolates included 129 individual *Arthrobacter* strains based on their partial *16S* rRNA sequences (94, 10, and 25 isolates from the tobacco endosphere, santhopine-treated soil, and nicotine-treated soil, respectively; Data Files S1). Fifty-four isolates were then randomly selected for phylogenetic characterization using their near-complete *16S* rRNA sequences (Data Files S2), which included strains from two other bacterial genera, *Paenarthrobacter* and *Pseudarthrobacter*, recently reclassified from the genus *Arthrobacter* based on their chemotaxonomic traits (*30, 31*) (Fig. S4). Given that these strains were found to be closely related to other *Arthrobacter* isolates at the *16S* rRNA gene sequence level as well as the whole-genome level (see below), these isolates were hereafter considered part of a single genus *Arthrobacter*. Although the isolates from santhopine- and nicotine-treated soils were relatively diverse in their *16S* rRNA sequences, most isolates from tobacco roots (27 of 30) were closely related to each other, forming a monophyletic clade with the *Arthrobacter nicotinovorans* DSM420, together with two isolates derived from santhopine-treated soil. These results indicate the predominance of a taxonomically limited range of *Arthrobacter* in the tobacco endosphere, which cannot be explained by the presence of santhopine or nicotine alone.

### Subspeciation of the genus *Arthrobacter* driven by a whole genome-scale rearrangement

We hypothesized that the specific enrichment of this particular subset of *Arthrobacter* isolates in the tobacco endosphere might mediated by whole genome-scale functional properties specific to these isolates. To test this, a taxonomically diverse set of *Arthrobacter* strains was selected from our culture collection (20 strains in total; Fig. S4) for whole genome sequencing. We obtained high-quality, closed genomic sequences (1–3 contigs, except for the strain NicSoilA12 with 12 contigs; Data Files S3), which were then compared to the genomes of 79 previously sequenced *Arthrobacter* isolates obtained from a variety of other hosts, such as *A. thaliana*, *Lotus japonicus*, and *Chlamydomonas reinhardtii*, as well as from soils and other environments (Data Files S4). First, their phylogenetic relationship was inferred based on the concatenated multiple sequence alignments of vertically inherited, single-copy genes (AMPHORA genes) (*32*). This analysis defined three distinct phylogroups within the genus *Arthrobacter*, hereafter referred to as sublineages A to C, respectively (Fig. 3). All isolates from tobacco roots and santhopine-treated soil (except for NtRootA9 and StoSoilB19) belonged to sublineage A, and the rest, including five strains from nicotine-treated soil, were classified into sublineage B. None of the newly sequenced isolates belonged to sublineage C. We classified each genome as plant-associated (“Plant”), soil-derived (“Soil”), or derived from other environment sources (“Other”) based on their origin of isolation (*15*). Sublineages A and B were overrepresented by the plant-associated isolates (19 of 31 and 22 of 43 “Plant” isolates, respectively). In contrast, sublineage C was mainly composed of isolates from other environments (16 of 25 “Other” isolates), such as animal, food, water, and extreme environment (Fig. 3; outer ring).

**Fig. 3.**
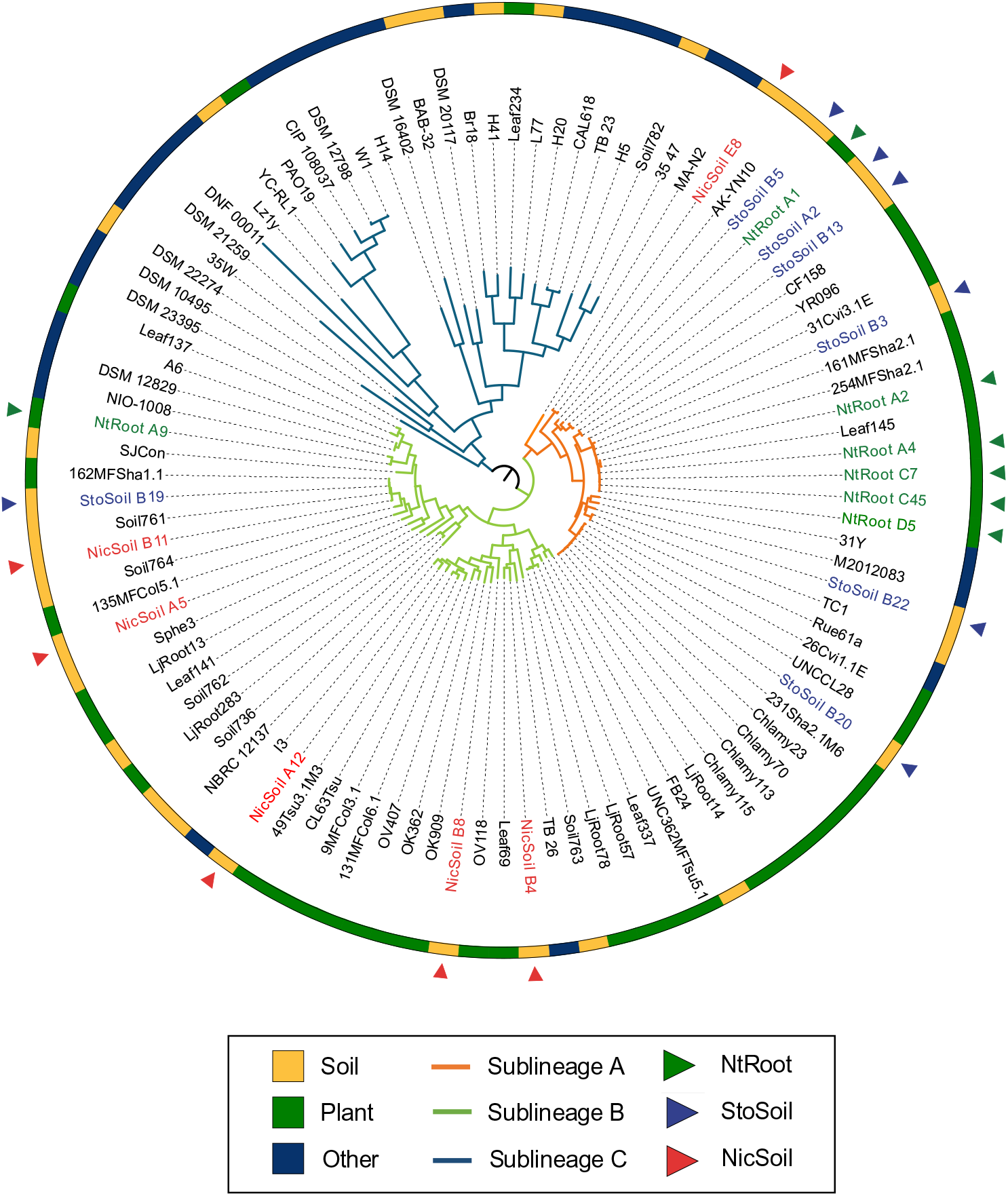
Whole genome-based phylogenomic reconstruction of the genus *Arthrobacter*. Phylogenetic tree of 99 *Arthrobacter* genomes inferred from aligned single-copy, vertically inherited marker genes using an MLE method. The branch colors represent the sublineages. The outer ring depicts the isolation source of each genome. Arrowheads indicate the newly sequenced isolates whose colors represent the origin of each strain.

To assess the functional potential of these sequenced strains, 99 representative genomes were annotated using the Kyoto Encyclopedia of Genes and Genomes (KEGG) database, resulting in 3803 KEGG Orthology groups (KOs). We also employed a *de novo* orthology prediction algorithm (*33*) and obtained 12,020 orthologous groups (OGs), the majority of which had no assigned functional prediction. Principal coordinates analysis (PCoA) was then performed using whole genome-level functional distances based on the presence or absence of OGs or KOs (*34*). This analysis showed that the functional distances based on *de novo* OGs, including genes without KEGG functional annotations, clearly explained the speciation of the sublineages defined by their AMPHORA phylogeny (Fig. 4A). In contrast, the functional distances based on the annotated KOs resulted in a single large cluster of genomes and did not correlate with the subspeciation (Fig. 4A), suggesting that the diversification of these lineages is found primarily in the non-annotated portion of their pan-genome. Strain MA-N2 belonged to sublineage A based on the phylogenetic analysis (Fig. 3), whereas it was clustered with sublineage B isolates in the PCoA plot of the genomic composition (Fig. 4A), which may be explained by an intermediate status of this strain between sublineages A and B. Based on its encoded functional capabilities, we decided to classify this strain as a constituent of sublineage B for the subsequent genomic analyses. Notably, the number of ORFs and OGs predicted in the genomes from sublineage A was significantly higher than those from sublineages B and C. In contrast, those from sublineage C appeared to have the smallest genome size (Fig. 4B), while the number of KOs was comparable among sublineages. The larger genome size of soil-derived and/or plant-associated bacteria than non-plant-associated bacteria was also reported in a previous study comparing a taxonomically broader set of bacterial genomes from a variety of sources (*15*). Overall, these results suggest that the subspeciation of the *Arthrobacter* genus is linked to adaptation to different environments and might have been associated with gains of genes in sublineages A and B and/or losses in sublineage C, whose functional characteristics are yet underexplored.

**Fig. 4.**
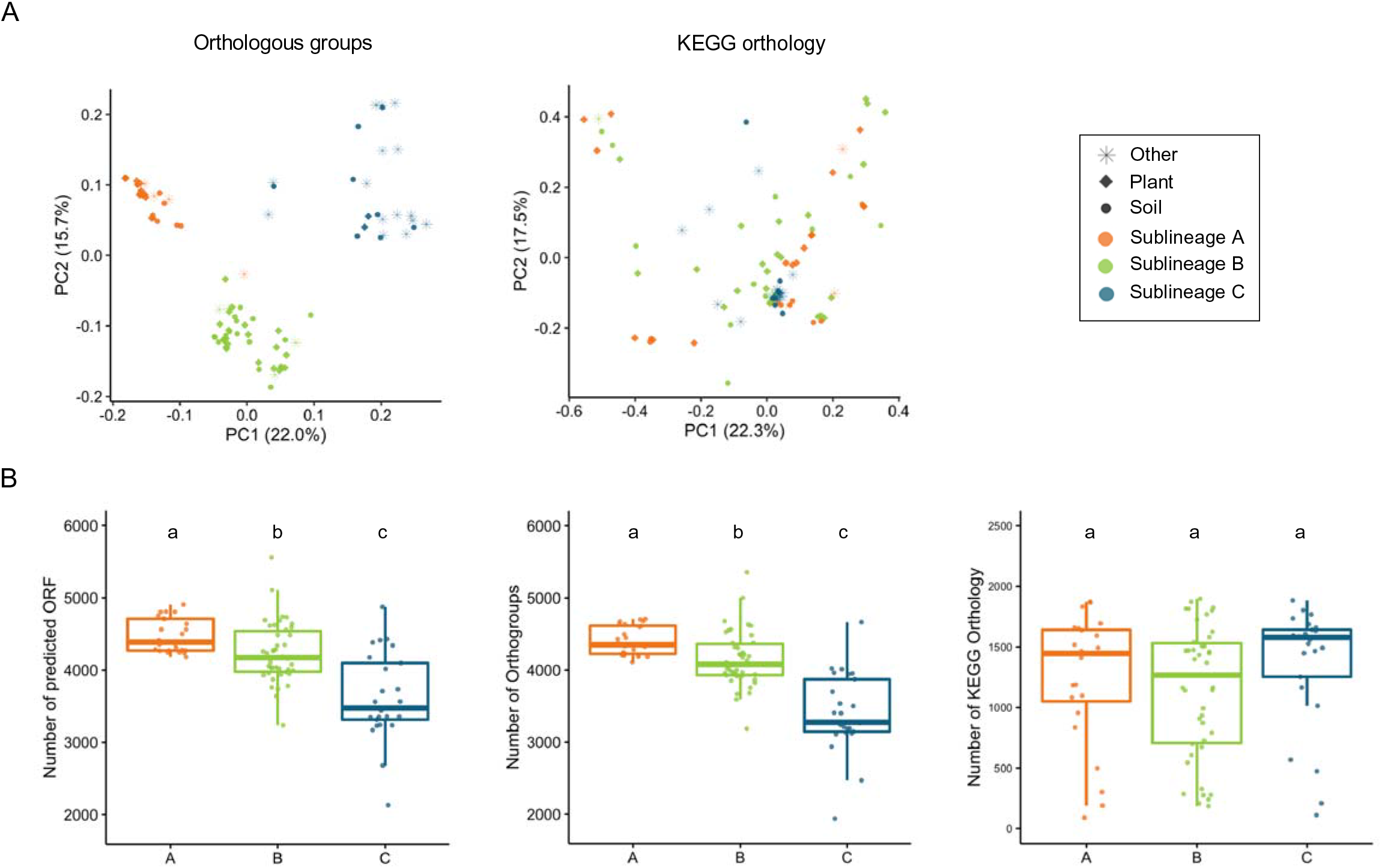
Functional diversity analysis and genomic size comparison between *Arthrobacter* genomes. **(A)** PCoA plots projecting functional distances between 99 *Arthrobacter* genomes based on the predicted OGs and the KO. Each point represents each genome. Shapes correspond to the isolation sources, and colors represent the sublineages defined by their AMPHORA phylogeny. **(B)** Genome size comparison of each *Arthrobacter* sublineage based on the number of predicted ORFs, OGs, and KOs. Different letters indicate statistical difference corresponding to a pairwise Wilcoxon’s rank-sum test (*P* < 0.05).

### Distinctive physiological properties between *Arthrobacter* sublineages A and B

To examine the sublineage composition of *Arthrobacter* in microbial communities in both the tobacco endosphere and santhopine or nicotine-treated soils, we analyzed sequences of the ASVs belonging to the family Micrococcaceae and the V4 region sequences of the *16S* rRNA genes in our 20 genome-sequenced *Arthrobacter* strains. This analysis revealed that ASV01 and ASV02 represented the *Arthrobacter* sublineages A and B, respectively (Fig. S5A). ASV02 was predominant in all the samples analyzed (Fig. S5B), including the santhopine- and nicotine-treated soils as well as the tobacco endosphere. However, despite the identical growth conditions employed for isolation from all types of samples, strains isolated from tobacco roots almost exclusively belonged to sublineage A (ASV01), whereas those from santhopine- and nicotine-treated soils were widespread across both sublineages (Data Files S1). The inconsistency between culture-dependent and culture-independent datasets may be explained by the possibility that sublineage A isolates colonize endophytically while those from other sublineages colonize only epiphytically. This distinctive property may be accounted for by different metabolic activities before and/or after the isolation procedures, such that sublineage B isolates become dormant in the endosphere or severely damaged by the root tissue disruption. Nonetheless, these findings point to a clear distinctiveness of sublineages A and B in their physiological properties during interactions with tobacco roots.

### *Arthrobacter* sublineage A has unique genetic components

To assess whether the identified *Arthrobacter* sublineage A has genomic signatures linked with a plant-associated or endophytic lifestyle, we surveyed their genomes for the prevalence of genes previously identified as relevant for root colonization in well-characterized rhizobial endophytes (*35, 36*). Among 101 genes reported in rhizobia, 61 genes were identified at least in one of the genomes in the genus *Arthrobacter* (Fig. 5A; Data Files S5). The proportion of the genes relevant for root colonization found in the genomes from sublineages A and B was significantly higher than in the genomes from sublineage C, although it was not significantly different between sublineages A and B (Fig. 5B). A similar proportion of genes required for root colonization was also detected between plant- and soil-derived isolates, which was higher than that in environment-derived strains (“Other”; Fig. 5C). This revealed that sublineages A and B share certain genomic features associated with root colonization, which supports the idea that these sublineages are both better adapted to the plant niche but does not explain the physiological distinctiveness between these sublineages.

**Fig. 5.**
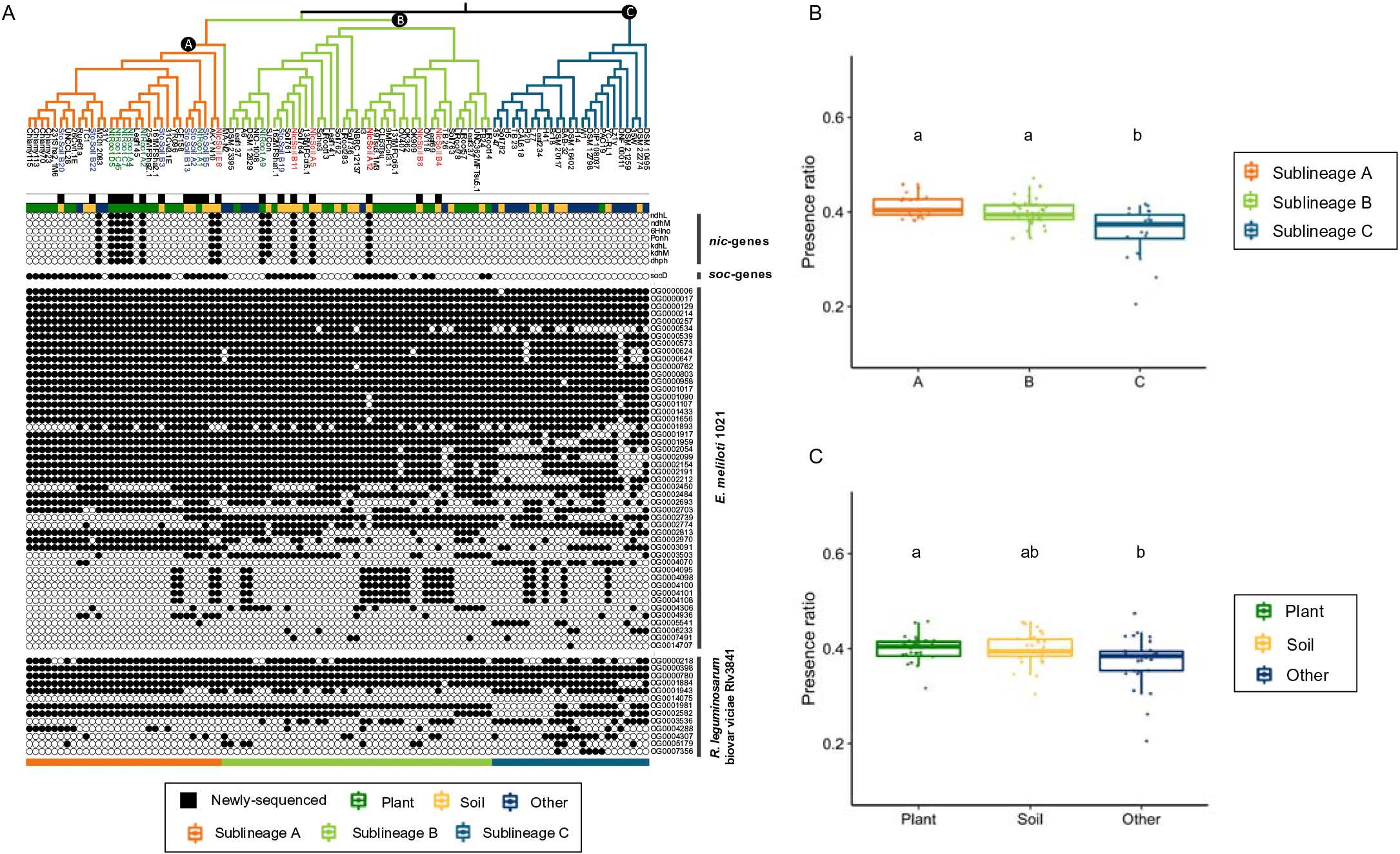
Phylogenetic distribution of *soc*, *nic*, and genes relevant for root colonization. **(A)** The tree indicates the AMPHORA phylogeny. The branch colors represent the sublineages. Text colors represent the origin of newly sequenced bacterial isolates. The first and second strips indicate the newly sequenced genomes and the isolation origins of each strain, respectively. Alphabets at node indicate the most recent common ancestor of each sublineage that was used for the ancestral character estimation. The presence and absence of *soc* and *nic* genes and the genes required for root colonization in rhizobial endophytes (*Ensifer meliloti* 1021 and *Rhizobium leguminosarum* biovar *viciae* Rlv3841) are represented by closed and opened circles, respectively. **(B** and **C)** For the genes needed for root colonization, the proportion of the number of genes present in each genome to the total number of genes present in at least one of these genomes (62 genes) was computed and shown as a boxplot, comparing between sublineages **(B)** and isolation origins **(C)**. Different letters indicate statistical difference corresponding to a pairwise Wilcoxon’s rank-sum test (*P* < 0.05).

To further disentangle the functional differences between sublineages A and B at the genome level, the ancestral characters of each *Arthrobacter* sublineage were estimated based on KO or OG profiles by a maximum likelihood estimation (MLE) approach. Using this analysis, we obtained a list of genes whose gains and losses were associated with the subspeciation events and identified 27 (KO) and 349 (OG) ancestral characters unique to sublineage A, and no ancestral Kos specific to sublineage B or C (Fig. S6; Data Files S6). Most ancestral KOs specific to sublineage A (27 KOs with 50 KEGG functional assignments) were functionally categorized as part of carbohydrate and amino acid metabolism (6 and 8 genes, respectively) and biosynthesis of bacterial secondary metabolites (5 genes). Among those genes specific to sublineage A, no genes known to be involved in root endophytic colonization, such as chemotaxis/bacterial motility, quorum sensing, and secretion systems (*37*), were identified. These findings illustrate that the *Arthrobacter* sublineage A possesses unique genomic traits that contribute to its distinctive biological properties compared to sublineage B.

### Endophytic colonization of *Arthrobacter* in tobacco roots is mediated by unique catabolic capacities

Although comparative genomics analyses revealed that the genes generally important for plant root colonization were widely conserved in *Arthrobacter* sublineages A and B, the enrichment of *Arthrobacter* in the root endosphere was observed only in tobacco and not in the other tested plant species (Fig. 2B). Moreover, most bacterial strains isolated from tobacco roots formed a monophyletic clade within sublineage A (Fig. S4). These data indicate additional genetic features that led to the tobacco-specific endophytic competence of these *Arthrobacter* isolates, which likely occurred after the divergence of sublineages A and B. To assess this, we surveyed the presence of the catabolic genes for santhopine and nicotine, both of which are characteristic of tobacco roots and induced the enrichment of *Arthrobacter* species by inoculation into the soils (Fig. 1C and D).

*soc* genes involved in santhopine catabolism were detected in all isolates derived from the tobacco endosphere and santhopine-treated soil, except for NtRootA9, although not necessarily in strains isolated from nicotine-treated soil (Fig. 5A; Fig. S7). Bacterial *nic* genes, which are involved in nicotine catabolism, in contrast, were frequently identified in isolates from the tobacco endosphere and nicotine-treated soil but not in isolates from santhopine-treated soil (Fig. 5A; Fig. S7). An *in vitro* growth assay confirmed that the isolates possessing *soc* and/or *nic* genes degraded representative metabolites, except for NtRootA9, which did not degrade nicotine despite the presence of *nic* genes in its genome (Fig. S7). These data revealed that the catabolic potential of these bacterial isolates largely correlated with the presence of representative metabolites in the environment from which they were isolated. Among the isolates derived from the tobacco endosphere, only those in sublineage A catabolized both metabolites. Importantly, the distribution of *soc* and *nic* gene sets in sublineages A and B was independent of host plant species of origin or of isolation site (Fig. 5A), indicating that the acquisition of these genes was not associated with niche establishment in the tobacco endosphere but rather with adaptation to these metabolites independently of the environments. Ancestral character estimation indicated that the most recent common ancestor of sublineage A already possessed the *soc* genes (*P* = 0.983), whereas *nic* genes were subsequently acquired during the divergence of the sublineages likely via HGT events (*P* = 0.980) (*38*). Collectively, our findings imply an evolutionary scenario according to which the presence of the *soc* genes, as well as other ancestral genes specifically found in sublineage A, predisposed ancestral *Arthrobacter* for colonization of the tobacco endosphere, possibly triggered by HGT of the *nic* genes.

## Discussion

We uncovered a link between the catabolic capacity toward santhopine and nicotine in *Arthrobacter* and the host species-specific enrichment of this genus in the tobacco endosphere. Although both santhopine and nicotine treatments resulted in an enrichment of *Arthrobacter* in soil bacterial communities, these metabolites differ in biological activities and their corresponding catabolic processes in bacteria. Santhopine is an Amadori compound (*39*) composed of glucose and glutamine and requires only a few enzymatic steps to be utilized in bacteria (*40, 41*). The presence of *soc* genes responsible for the catabolism of a wide range of Amadori compounds is limited to a set of bacterial families, such as Rhizobiaceae, Microbacteriaceae, and Micrococcaceae (*24*), rendering santhopine a nutrient source for specific groups of bacteria. It has been reported that *Salmonella*, a foodborne animal pathogen, has a unique ability to utilize fructose-asparagine, also an Amadori compound, as a carbon and nitrogen source to eliminate potential competitors in the inflamed intestine (*42*). We speculate that santhopine provides *Arthrobacter* strains harboring *soc* genes a competitive advantage against other bacteria in the tobacco endosphere. Soc enzymes can metabolize a wide range of Amadori compounds (*39–41*), and Amadori compounds occur in various environments, such as foods, decomposing plants, human blood, and plant tissues (*32, 43, 44*). Thus, it is likely that soil-inhabiting *Arthrobacter* species have originally acquired *soc* genes to utilize soil-borne Amadori compounds as their nutrients.

In contrast, nicotine exhibits antimicrobial activity against various bacteria and fungi (*45*), and a series of enzymatic reactions via the *nic* gene cluster (located on the pAO1 megaplasmid) are required for its catabolism (*38*). Thus, detoxification of nicotine by *nic* genes might be beneficial for colonization of the tobacco endosphere. In addition, *Arthrobacter* AK-YN10 and SJcon strains harboring *nic* genes have been isolated from agricultural soils contaminated by nitrogen-containing pesticides (atrazine and 2-chloro-4-nitrophenol, respectively) (*46, 47*), pointing to the contribution of *nic* genes to catabolize not only nicotine but also other alkaloid-like compounds. Members of the *Arthrobacter* genus have often been isolated from contaminated soils with benzene derivatives, polycyclic aromatic compounds, and nitrogen-containing pesticides (*48*), indicating that *nic* genes are generally associated with adaptation to soil environments rich in alkaloids or alkaloid-like compounds, including nicotine.

The importance of santhopine and nicotine for the tobacco-specific endophytic competence of the genus *Arthrobacter* raises the question of how these bacteria might have acquired the ability to catabolize these PSMs and whether this metabolic capacity can alone explain the observed host-specific interactions. Among the three distinctive *Arthrobacter* sublineages defined in this study, sublineages A and B appeared to be better adapted to the soil and/or plant environments. *Arthrobacter* strains isolated from wild tobacco (*Nicotiana attenuata*) and holm-oak exerting plant growth-promoting activity were also grouped with sublineages A and B isolates based on their *16S* rRNA sequences (Fig. S8) (*49, 50*). Our genomic analyses revealed the presence of functional properties enriched in sublineage A, including the *soc* genes, which are possibly required for the interaction with tobacco, in addition to those widespread across sublineages A and B for the plant-associated lifestyle. Notably, isolates that exhibited the *in vitro* activity to catabolize both santhopine and nicotine (Fig. S7) and the isolates from soils contaminated by alkaloid-like compounds possessing both *soc* and *nic* genes (AK-YN10 and SJcon) were found in both sublineages A and B. In contrast, almost all isolates derived from tobacco roots and the isolate from tobacco waste (M2012083) were specifically found in sublineage A. This finding suggests that other genes enriched in sublineage A, besides the *soc* genes, also play a role in their ability to colonize the tobacco endosphere abundantly. Overall, our results indicate that the horizontal acquisition in *Arthrobacter* of the *nic* genes to catabolize nicotine (*38*) and the *mas2* gene to synthesize santhopine in tobacco (*16*) mediates interactions between this bacterial genus and tobacco plants (Fig. 6). The observation that isolates from tobacco roots and the isolate from tobacco waste were taxonomically distant from each other within sublineage A (Fig. 3), suggests independent acquisition events of the *nic* operon. Thus, the observed interactions between *Arthrobacter* and tobacco may not be the consequence of co-evolution between the two partners, but rather the result of a process of ecological fitting.

**Fig. 6.**
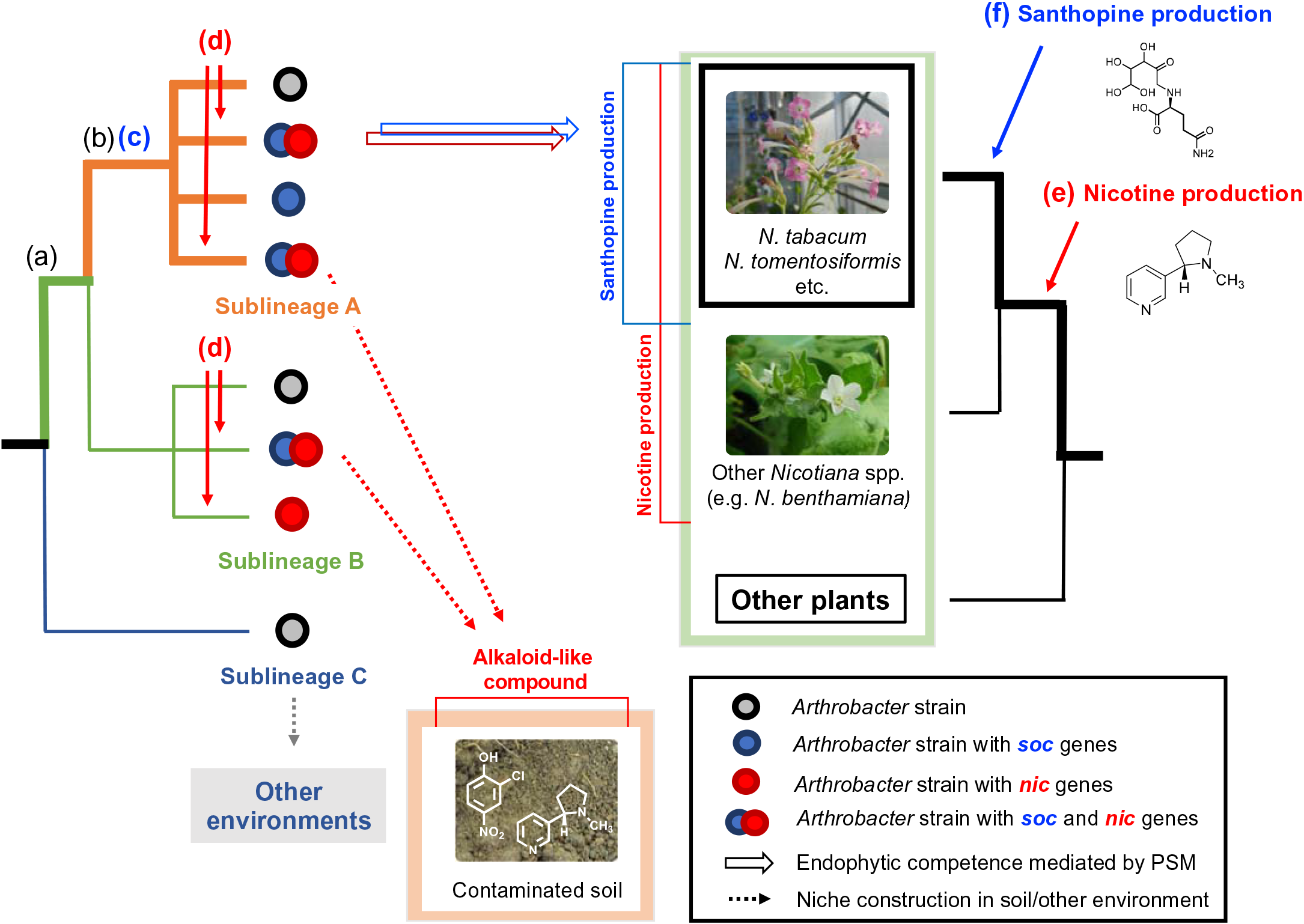
Proposed evolutionary model of a host-specific root microbiota assembly mediated by bacterial catabolism of a combination of PSMs synthesized by the host plant. Anticipated acquisition events of plant and bacterial characteristics related to the interaction between tobacco and *Arthrobacter* are represented by alphabets. (a) Acquisition of genes required for plant-associated lifestyle, which are conserved in sublineages A and B. (b) Acquisition of genes specifically enriched in sublineage A. (c) Acquisition of *soc* genes. (d) HGT events of *nic* genes, which have been frequently occurred within sublineages A and B. (e) Acquisition of genes required for nicotine production. (f) HGT event of the *mas2* gene for santhopine production. (b) and (c) may originally be associated with adaptation to the soil environments. The co-occurrence of (a) to (d) was observed only in the isolates from tobacco roots. The host-specific enrichment of *Arthrobacter* in tobacco roots was possibly triggered by (d) and (f) in bacteria and plants, respectively.

Interestingly, *Arthrobacter* strains isolated from tobacco were able to catabolize both santhopine and nicotine, although these two compounds belong to unrelated classes of PSMs (Fig. 1A). Similarly, both compounds mediate the enrichment of the same bacterial genus in the soil bacterial community. Therefore, the capacity of the host to produce a combination of both compounds mediates the enrichment of *Arthrobacter* in the root compartment and influences root microbiota composition. This finding contrasts with the effect of PSMs in soybean, isoflavones, and soyasaponins, which enriched different bacterial families in soils treated with these metabolites (*13, 14*). Considering the fact that a plant produces a wide range of lineage-specific PSMs (*3*), it is likely that host-specific microbiota assembly is partially achieved by the joint action of multiple PSMs. This mechanism might explain why the disruption of a single metabolic pathway in plants typically results in only minor changes in the root microbiota composition (*10–12*). Together with previous studies showing the impact of PSMs on the root-associated community structure (*10–12*), our findings suggest a mechanistic model of host-specific root microbiota assembly mediated by the catabolic potential of bacteria toward a cocktail of host-specific PSMs.

## Materials and methods

### Chemicals and soil

Santhopine was synthesized as described previously (*16*). Glucose (1.35 g) was dissolved in 15 mL AcOH at 95°C, and then L-glutamine (300 mg) was added. The reaction mixture was stirred about 3 min at 95°C. Five times volume of ethyl acetate was added to the reaction mixture and centrifuged at 4000×*g* for 10 min. Then, the precipitate was dried completely with an evaporator and dissolved in 10 mL water. This sample was simply purified by a Sep-Pak C18 20 cc Vac cartridge (Waters, Milford, MA, USA). The flow-through fraction containing santhopine was freeze-dried and dissolved in methanol. Dissolved samples were purified by preparative high-performance liquid chromatography (HPLC) using COSMOSIL Sugar-D packed column (10 mm I.D. × 250 mm; Nacalai Tesque, Kyoto, Japan). Santhopine was eluted at 4.0 mL min^−1^ flow rate with a mobile phase consisted of (A) water and (B) acetonitrile. The stepwise gradient program was as follows: 95% A, 0 to 5 min; isocratic at 50% A, 5 to 15 min. Absorbance at 210 nm was monitored. Fractions containing santhopine were pooled and completely dried by an evaporator and freeze-dried.

The structure of the synthesized sample was confirmed by liquid chromatography-mass spectrometry (LC-MS) analysis and nuclear magnetic resonance (NMR) spectroscopies (^1^H and ^13^C). LC-MS analysis was performed using an ACQUITY UPLC H-Class system (Waters) with an ACQUITY UPLC BEH amide column (1.7 μm, 2.1 × 100 mm; Waters) at 40°C and an SQ Detector 2 (Waters). The injection volume of each sample was 2 μL and the flow rate was 0.2 ml min^−1^. The LC mobile phase consisted of (A) water, (B) acetonitrile, and (C) water containing 4% (v/v) formic acid. The elution program was a linear gradient of 90 to 50% A supplied with 5% C at 0 to 15 min. Santhopine was analyzed in a positive electrospray ionization with the following conditions: cone voltage, 25 V; capillary voltage, 3.15 kV; source temperature, 150 °C; desolvation gas temperature, 400°C; and nebulizer and desolvation N_2_ gas flow rates, 50 and 800 L h^−1^, respectively. Santhopine was detected using selected ion recording (SIR) mode with *m/z* 309.2 and multiple reaction monitoring (MRM) mode with the following conditions: transition, *m/z* 309.2 > 130.1; cone voltage, 25 V; and collision energy, 20 eV. Data acquisition and analysis were performed using MassLynx 4.1 software (Waters).

^1^H and ^13^C NMR spectra were measured at 25°C on a 400 MHz and 100 MHz instrument in deuterated water, respectively. ^1^H NMR (400 MHz, D_2_O, 25 °C): δ 3.92–3.89 (m, 2H, H(3′), H(6′)), 3.76 (dd, 1H, H(4′)), 3.66–3.59 (m, 3H, H(3′), H(6′), CH(4)), 3.20–3.18 (m, 2H, CH2(1′)), 2.40–2.38 (m, 2H, CH2(2)), 2.01–1.99 (m, 2H, CH2(3)). ^13^CNMR (100 MHz, D2O, 25 °C) : δ 177.8 (C=O(C-1)), 172.8 (C=O(C-5)), 95.3 (C-2′), 69.9 (C-4′), 69.3 (C-5′), 68.9 (C-3′), 62.6 (CH(C-4)), 63.9 (C-6′), 52.6 (CH2(C-1′)), 31.1 (CH2(C-2)), 24.9 (CH2(C-3)) (Fig. S9; Table S1). Chemicals were obtained from Wako Pure Chemical Industries (Osaka, Japan) or Nacalai Tesque, unless otherwise stated. Field soil was collected from a field at the Kyoto University of Advanced Science (KUAS), Kameoka, Kyoto, Japan (34°99′38′′N, 135°55′14′′E), as described previously (*13*).

### Treatment of field soil with santhopine and nicotine

Three different concentrations of santhopine or nicotine solution (low, 50 nmol g^−1^ soil; middle, 250 nmol g^−1^ soil; high, 1000 nmol g^−1^ soil) were applied every 3 days for five times, as described previously (*13*). Sterile water was applied as a control. Tubes were incubated at room temperature in the dark. The concentrations of santhopine or nicotine solution were adjusted every time to keep the soil water content ratio at 30%. After incubation, total DNA was extracted using DNeasy PowerSoil Kit (Qiagen, Hilden, Germany) according to the manufacturer’s protocol. Extracted DNA was quantified using Qubit Quantification Platform dsDNA HS Assay Kit (Invitrogen, Carlsbad, CA, USA) and stored at −80°C.

### Sample collection from tobacco roots

Seeds of *N. tabacum* cv. Burley 21 (tobacco) was provided by Japan Tobacco, Inc. (Tokyo, Japan). Tobacco seeds were surface sterilized with 70% ethanol (EtOH) for 1 min and 1% sodium hypochlorite (NaClO) for 10 min and rinsed five times with sterile water. Plants were germinated on Murashige and Skoog medium (Wako Pure Chemical Industries) supplemented with 0.8% agar. Plants were grown for 2 weeks in a cultivation room set at 28°C under a light/dark (16/8 h) photoperiod. Tobacco seedlings were then transferred to plastic pots filled with soil collected at the KUAS and grown from 12 weeks in a greenhouse. The plants were fertilized every week using Hyponex (Hyponex Japan, Osaka, Japan).

The rhizosphere soil was collected using phosphate-buffered saline (PBS) after removing the loosely attached soil on the roots removed by gentle shaking, as described previously (*6*), with some modification. Roots were pooled into a 1000 mL glass beaker filled with 500 mL sterile PBS containing 130 mM NaCl, 7 mM Na_2_HPO_4_, 3 mM NaH_2_PO_4_ (pH 7.0), and 0.02% Silwet L-77, and washed on a shaking platform for 5 min at 100 rpm. After removing the roots, the buffer was centrifuged (5000×*g*, 10 min), and the resulting pellet was defined as the rhizosphere soil. Then, roots were placed in a new PBS and sonicated for 5 min. The resulting pellet collected by the centrifugation (5000×*g*, 10 min) was defined as the rhizoplane soil. After washing with tap water, the roots were surface sterilized with 70% EtOH for 1 min and 1% NaClO. The surface-sterilized roots were then washed five times with sterile water and stored −80°C until DNA extraction. The remaining root tissues were immediately subjected to bacterial isolation.

### Sample collection from field-growth plants

Tomato seeds (cv. Beni-Suzume) were sown in a culture soil, a 1:1 mixture of vermiculite and Tsuchitaro (Sumitomo Forestry Landscaping, Tokyo, Japan), and grown for 13 days at 28°C under a under light/dark (16/8 h) photoperiod. The seedlings were then planted in the field on May 9, 2019. Seeds of soybean (cv. Enrei) and bitter melon were directly sown in the field on May 31, 2019. All plants were sampled on August 1, 2019. After collecting rhizosphere soils as described above, the roots washed with tap water were subjected to DNA extraction as an endosphere compartment. Soybean and tomato roots for DNA extraction were collected from five and two plants per sample, respectively.

### Bacterial community analysis using *16S* rRNA amplicon sequencing

For the amplicon sequencing of the *16S* rRNA gene, total DNA was extracted from the soil and plant samples using DNeasy PowerSoil Kit (Qiagen) according to the manufacturer’s protocol. Extracted DNA was quantified using Qubit Quantification Platform dsDNA HS Assay Kit (Invitrogen). PCR amplification of the region V4 of *16S* rRNA was performed with KOD Fx Neo Polymerase using first PCR primers 515F (5′-<u>ACACTCTTTCCCTACACGACGCTCTTCCGATCT</u>-GTGCCAGCMGCCGCGGTAA-3′) and 806R (5′-<u>GTGACTGGAGTTCAGACGTGTGCTCTTCCGATCT</u>-GGACTACHVGGGTWTCTAAT-3′) consisted of the Illumina (San Diego, CA, USA) paired-end adapter sequences (underlined) and *16S* rRNA gene-specific sequences. The first PCR mixture for the soil DNA amplification consisted of 10 ng DNA template, 12.5 μL of 2× reaction buffer, 5 μL of 10 mM dNTP, 0.75 μL of 10 μM primers (each), 0.25 μL of KOD Fx Neo (Toyobo, Osaka, Japan), and 0.75 μL of nuclease and DNA-free water. The thermal programs of the first PCR for soil samples were as follows: 2 min at 94°C, 20 cycles of 10 s at 98°C, 30 s at 50°C, and 30 s at 68°C. For the first amplification of the 25 μM peptide nucleic acids were applied: mPNA (N-term-GGCAAGTGTTCTTCGGA-C-term) and pPNA (N-term-GGCTCAACCCTGGACAG-C-term), respectively (Panagene Inc., Daejeon, South Korea), to block the amplification of contaminating sequences from a eukaryotic host. PCR reaction was conducted using the following thermal programs: 2 min at 94°C, 25 cycles of 10 s at 98°C, 10 s at 78°C, 30 s at 50°C, and 30 s at 68°C. The amplified fragments were purified using Ampure magnetic beads (Beckman Coulter, Danvers, MA, USA) according to the manufacturer’s protocol. The second PCR mixture content was the same as the first PCR mixture for the soil DNA using primers provided from FASMAC Co., Ltd. (Kanagawa, Japan) and amplified using the following thermal conditions: 2 min at 94°C, 9 to 10 cycles of 10 s at 98°C, 30 s at 50°C, and 30 s at 68°C. The purification and quantification of PCR products were performed as described above. The PCR product mixture was sent to FASMAC to acquire 2 × 250 bp paired-end sequences using the MiSeq platform (Illumina).

Sequence data for the amplicons were analyzed using the QIIME2 platform, version 2019.04 (*51*). For all paired reads, the first 20 bases of both sequences were trimmed (to remove primer sequences), and the bases after 220 were truncated (to remove low-quality sequence data). Potential amplicon sequencing errors were corrected using DADA2 to produce an ASV dataset (*52*). Obtained ASVs were aligned using MAFFT (*52*), and a phylogenetic tree was contracted using FastTree 2 (*53*). Each ASV was assigned using a naïve Bayes classifier from the Greengenes release 13_8 dataset (*51*), and then the reads for chloroplasts or mitochondria were removed. The calculation of α and UniFrac distances, PCoA analysis for β-diversity, and Adonis PERMANOVA test for β group significance was performed using the QIIME2 platform. The Wilcoxon’s rank-sum test comparing weighted UniFrac distance was performed using R software. Statistical analyses of differentially abundant families and orders were performed using the edgeR library (*54*) by fitting a negative binomial generalized linearized model to the ASVs. The sequence dataset supporting the results of this study was submitted to the DNA Data Bank of Japan (https://www.ddbj.nig.ac.jp) and to be publicly available.

### Isolation of root- and soil-derived *Arthrobacter* and phylogenic analysis

Bacterial strains were isolated from the surface-sterilized roots of tobacco and the soils treated with santhopine or nicotine at 1000 nmol g^−1^ soil. The surface-sterilized roots were sectioned into smaller fragments and then homogenized with mortar and pestle in PBS. The santhopine/nicotine-treated soil was suspended in distilled water at 0.5 g soil mL^−1^. Homogenates and soil suspensions were diluted and distributed onto isolation media (for isolation from tobacco roots: M408, M715, TYG, and YEM; for isolation from chemical-treated soils: M408 and M715), as described by Bai et al. 2015 (*34*) (Data Files S1). Mineral salt buffer (MS) medium (*55*) with 1 mg mL^−1^ nicotine solution was also used for isolation from nicotine-treated soil (Data Files S1). Plates were incubated up to 7 days at 28°C. Colonies were picked from plates and sub-cultured on growth medium (10 g L^−1^ pepton, 10 g L^−1^ beef extract, and 5 g L^−1^ NaCl) and then preserved in 25% glycerol solution at −80 °C. Genomic DNA was extracted by the hot-alkaline DNA extraction method using the extraction buffer (25 mM NaOH, 0.2 mM EDTA, 40 mM Tris-HCl; pH 6.8). The *16S* rRNA genes were amplified using 10F (5′-GTTTGATCCTGGCTCA-3′) and 1500R (5′-TACCTTGTTACGACTT-3′) primers sets. PCR products were purified with Wizard Genomic DNA Purification Kit (Promega, Madison, WI, USA) according to the manufacturer’s protocol and directly sequenced using 10F primer to identify individual bacteria isolates at the genus level. Fifty-four bacterial isolates assigned as genus *Arthrobacter* from the bacterial culture collection for further phylogenic analysis (30, 10, and 14 individual *Arthrobacter* strains were isolated from the tobacco endosphere, and santhopine-treated soil, and nicotine-treated soil, respectively) were then randomly selected and sequenced using the 1500R primer. The resulting near-complete *16S* rRNA sequences (1202 bp) were aligned with the Clustal Omega 1.2.4 (*56*), and the resultant alignment was trimmed using trimal 1.4 (*57*). The *16S* rRNA sequence data for the reference strains were retrieved from the GenBank database by BLAST search. Phylogenetic trees were constructed by the MLE method using FastTree version 2.1 (*53*).

### Whole genome sequencing of *Arthrobacter* isolates

The genomic DNA of *Arthrobacter* isolates was extracted as described by Hahn and Hennecke with a modification (*58*). Briefly, the cells were lysed by the addition of 20 mg lysozyme, 10% sodium dodecyl sulfate, and proteinase K (20 mg mL^−1^) and incubated for 1 h at 37°C. The cell lysate was forced through a syringe (21G×1-1/2″ RB; Terumo Corporation, Tokyo, Japan) and extracted with phenol and chloroform. The genomic DNA was dissolved in TE buffer and kept at 4°C. For SMRTbell library preparation, each genomic DNA was fragmented at 20 kbp using a Megaruptor2 [Diagenode, Seraing (Ougrée), Belgium], and the library was constructed using SMRTbell Express Template Prep Kit 2.0 according to the manufacturer’s protocol (Pacific Biosciences, Menlo Park, CA, USA). The barcodes were attached to each fragmented genome, and the samples were pooled and cut off at 15 kbp using the BluePippin size selection system (Sage Science, Cummings Center Beverly, MA, USA). The genomic library was sequenced on a single PacBio sequel II system 2.0 cell. Genomes were assembled with HGAP4 via SMRTlink (version 8.0.0) using the specified genome sizes. The sequence dataset was submitted to the DNA Data Bank of Japan (https://www.ddbj.nig.ac.jp) to be publicly available.

### Whole genome sequencing of *Arthrobacter* isolates from *L. japonicus* and *C. reinhardtii*

Genomic DNA of bacterial strains isolated from *L. japonicus* rhizosphere and *C. reinhardtii* phycosphere samples was extracted and purified as described previously (*34*) and sequenced using the Illumina HiSeq 2500 at the Max Planck Genome Center (Cologne, Germany) with an insert size of 350 bp and an approximate depth of 5M reads per strain. Sequences were quality filtered by passing short reads through a quality and length-trimming filter using Trimmomatic (*59*) with the default parameters and subsequently assembling them using the A5 pipeline (*60*).

### Genome annotation and orthology inference

Public genome sequences of *Arthrobacter* were retrieved from the Integrated Microbial Genomes & Microbiomes system (https://img.jgi.doe.gov) and the National Center for Biotechnology Information (NCBI; https://www.ncbi.nlm.nih.gov). Using a total of 99 genomes, putative protein-coding sequences were predicted using Prokka (*61*). The annotation of candidate ORFs was then conducted using the KO database (*62*) by building a blast database using a de-replicated set representative sequences for each KO which was subsequently added to the Prokka pipeline as a custom Bacterial database. *De novo* orthology prediction was performed using OrthoFinder2 to obtain OGs (*33*).

### Comparative genomics and ancestral character reconstruction

For high-resolution phylogenetic inference of genomes, well-conserved (present in all genomes), single-copy bacterial genes were extracted using AMPHORA (*32*). The obtained sequences were aligned independently for each gene by Clustal Omega 1.2.4. Aligned sequences were trimmed and concatenated to infer a phylogenetic tree by an MLE method using FastTree version 2.1. PCoA of a functional diversity between sequenced isolates was performed, as described previously (*34*). Briefly, we generated a presence/absence profile of each KO and OG group was generated for each genome in the dataset. A dissimilarity matrix was generated based on Pearson’s correlation coefficients between each pair of genomes.

To avoid conflicts in the inference of ancestral characters, polytomies in the trees were resolved by inserting branches of zero length. Given the matrix of phyletic patterns based on KOs or OGs and the estimated species tree, an MLE approach (*63*) was used to estimate the ancestral state using the *ace* function implemented in the ape package (*64*) for R software. Ancestral characters of each *Arthrobacter* sublineage were defined as genes with likelihood values > 0.8 at a given node. The functions of identified OGs were further inferred by blasting representative amino acid sequences against the proteome dataset of the family Micrococcaceae at the NCBI (setting: *E*-value thread = 1 × 10^−50^).

To identify homologous *soc*, *nic*, and genes essential for the root colonization of rhizobia, deduced amino acid sequences of these genes were blasted against the proteome predicted by the *Arthrobacter* genome sequences (>50% query sequence coverage; *E*-value thread > 1 × 10^−5^). The presence of a santhopine catabolism operon was assessed by confirming the presence of *socD* from *Arthrobacter aurescens* TC1 (setting: *E*-value thread > 3 × 10^−100^; Table S2), as other *soc* components show high similarities to genes of amino acid metabolism genes, which might overestimate its prevalence. The presence of the *nic* gene cluster was surveyed using *nic* genes of *A. nicotinovorans* DSM420 as query sequences (setting: *E*-value thread = 0; Table S2). *ndhS* (CAD47953.1) and *kdhS* (CAD47946.1) genes were excluded due to their relatively short amino acid sequence lengths (156 and 160 amino acids, respectively). Phylogenetic trees and associated data were visualized using iTOL version 5 (https://itol.embl.de).

Statistical tests for the genome size and rhizobia gene proportions were performed using R software, according to the Wilcoxon’s rank-sum test. For statistical tests, we removed the genome sets of NtRootA4, C7, C45, D5, Chlamy70, 113, 115, and DSM_12829 were removed, as these genomes share more than 99% of their OG groups with NtRootA2, Chlamy23, and A6, respectively.

### Santhopine and nicotine degradation assay

Bacterial strains were cultured until strains reached the stationary phase at 28°C in the growth medium described above. Bacterial cells were collected and washed twice with MS medium. Bacterial pellets were resuspended in MS medium with 5 μg mL^-1^ santhopine or 1 mg mL^-1^ nicotine solution and incubated for 48 h. After centrifugation at 13,000×*g* for 5 min, supernatants were collected and stored in −80°C until use. The santhopine content in the supernatants was measured by LC-MS (Waters), as described below, with the modified LC condition. The LC mobile phase consisted of (A) water, (B) acetonitrile, (C) 2-propanol, and (D) water containing 4% (v/v) formic acid. The stepwise gradient program was isocratic at 70% B, 25% C, and 5% D, 0 to 3 min, and isocratic at 47.5% A, 35% B, 12.5% C, and 5% D, 3 to 8 min. Nicotine was analyzed by HPLC, as described by Häkkinen et al. 2004 (*65*).

## Acknowledgments

We thank Drs. Ryosuke Munakata, Takuji Ichino, and Shingo Miyauchi for their useful comments on this manuscript, Dr. Hisabumi Takase (Kyoto University of Advanced Science) for assistance with soil sampling, Dr. Masaru Nakayasu (Kyoto University) for the sampling of the field-growth plant, Japan Tobacco, Inc., for providing the tobacco seeds, and DASH/FBAS, the Research Institute for Sustainable Humanosphere, Kyoto University, for supporting the institutional setting. Some experimental measurements were carried out using the JEOL 400 MHz NMR spectrometer at the Institute for Chemical Research, Kyoto University. We also thank Enago (www.enago.jp) for English language review.

## Author contributions

T.S., R.T.N., and A.Sugiyama conceived and designed the experiments. T.S. and T.K. performed the synthesis of santhopine. T.S., Y.A., and R.T.N. performed the data analysis for the bacterial community. T.S. performed the isolation and characterization of bacterial strains. T.S., S.M., A.Shibata, W.S., and K.S. performed whole genome sequencing. T.S, R.G.-O., and R.T.N. performed comparative genomics analysis. K.Y., R.T.N., and A.Sugiyama supervised the entire study. T.S., R.T.N., and A.Sugiyama wrote the manuscript with the contribution of the other authors.

## Data and materials availability

All raw data and scripts as well as intermediate data are available at https://www.mpipz.mpg.de/R_scripts. Additional data related to this paper will be made available upon request.

## Supplementary materials

**Fig. S1.**
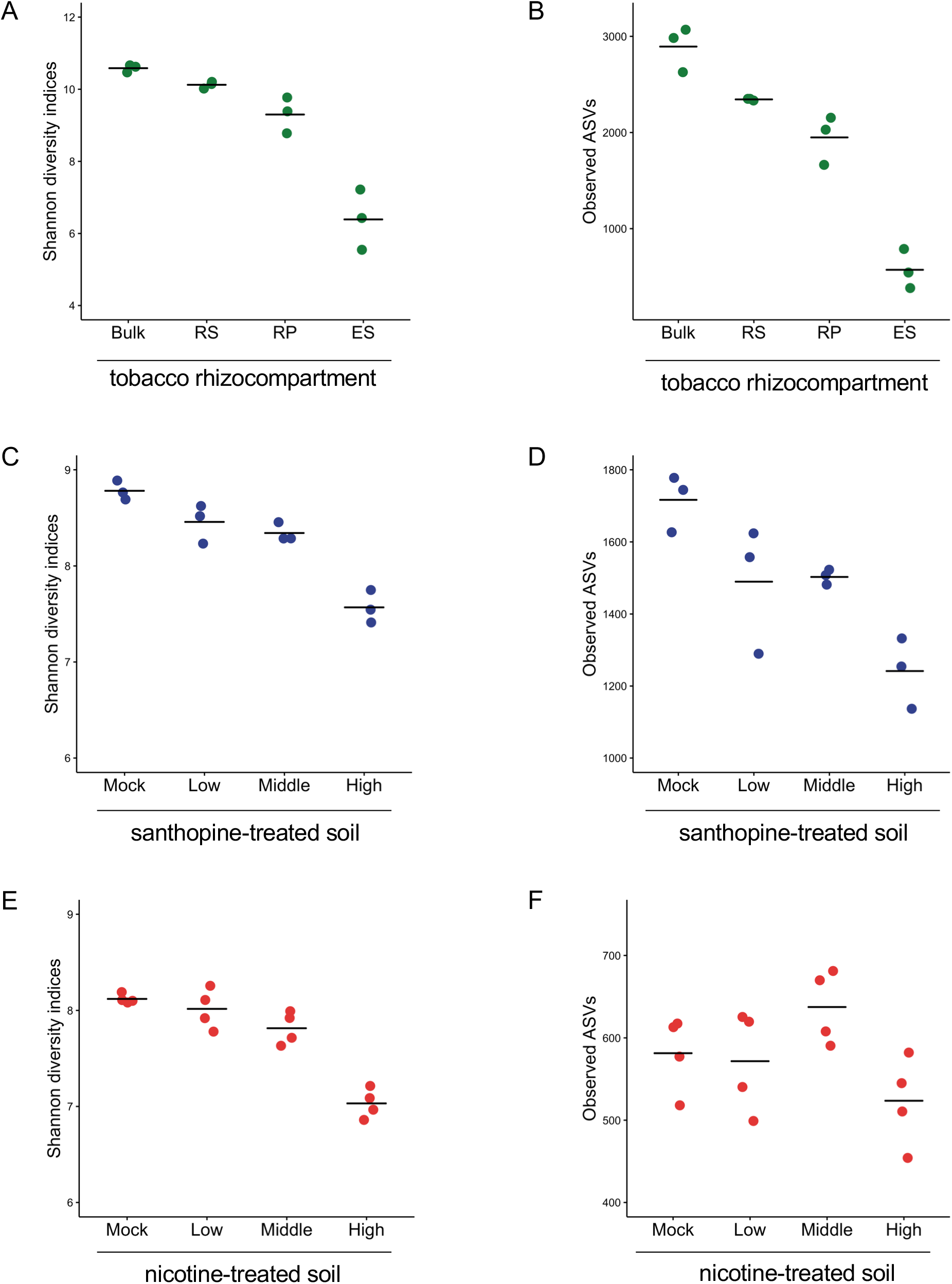
α-diversity of bacterial communities of each sample. Number of observed ASV index (**A, C, and E**) and Shannon diversity index **(B, D, and F)** of each sample are shown as individual points. Horizontal bars represent mean within a sample.

**Fig. S2.**
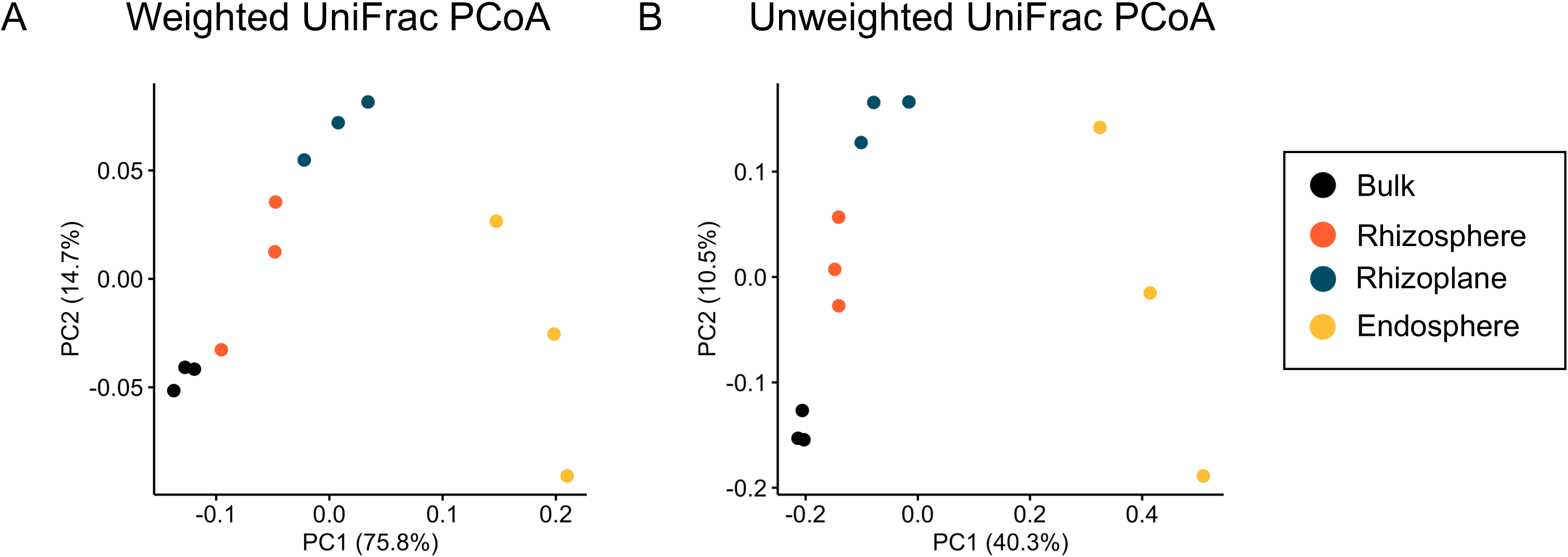
PCoA of bacterial communities of tobacco roots. PCoA based on **(A)** weighted and **(B)** unweighted UniFrac distances.

**Fig. S3.**
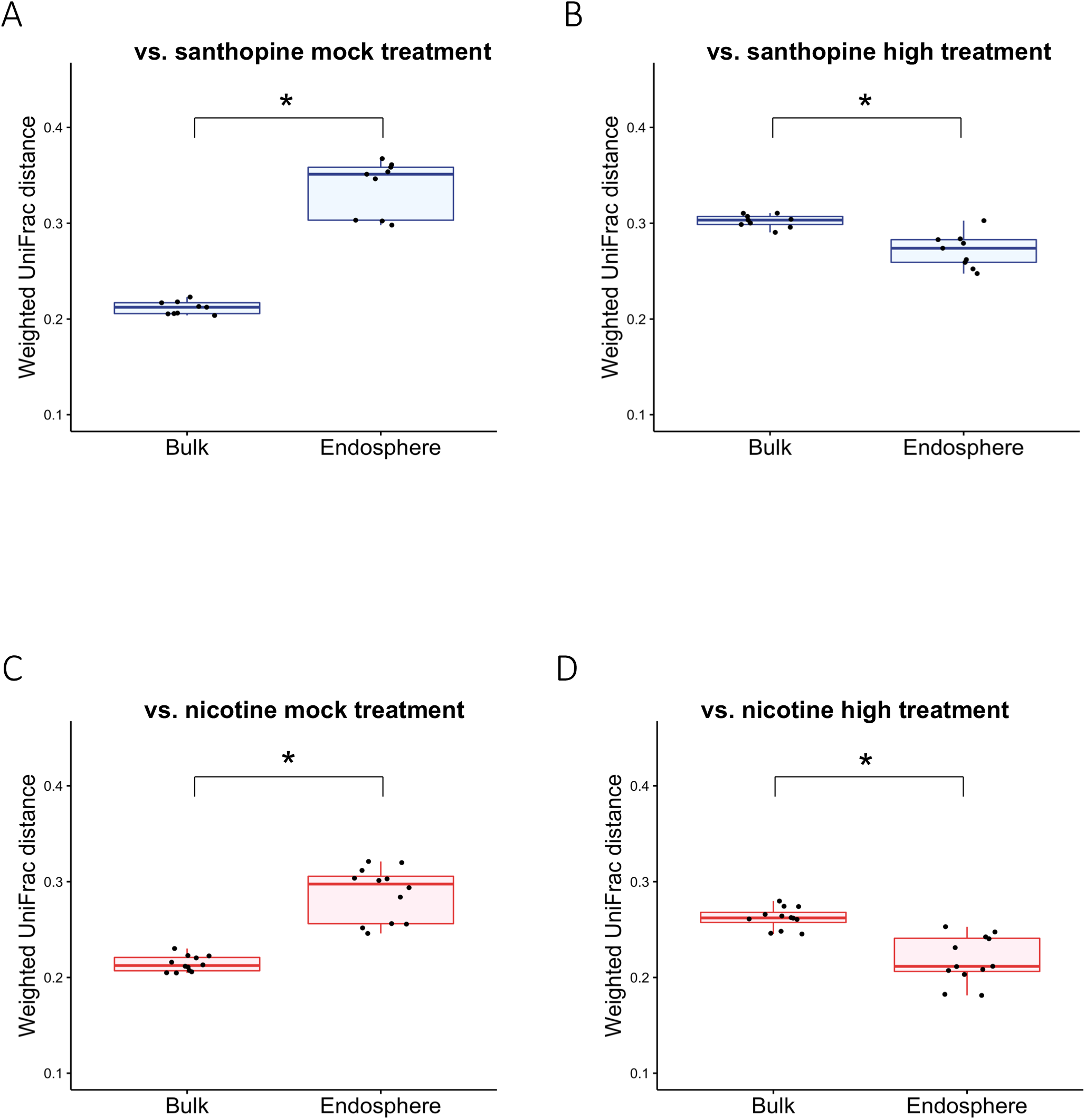
Comparison of bacterial communities in santhopine- and nicotine-treated soils to those of the tobacco endosphere. Weighted UniFrac distance between santhopine mock treatment and bulk/endosphere of tobacco **(A)**, between santhopine treatment at the highest concentration and bulk/endosphere of tobacco **(B)**, between nicotine mock treatment and bulk/endosphere of tobacco **(C)**, and between nicotine treatment at the highest concentration and bulk/endosphere of tobacco **(D)**. Asterisks indicate statistical difference corresponding to a Wilcoxon’s rank-sum test (*P* < 0.05).

**Fig. S4.**
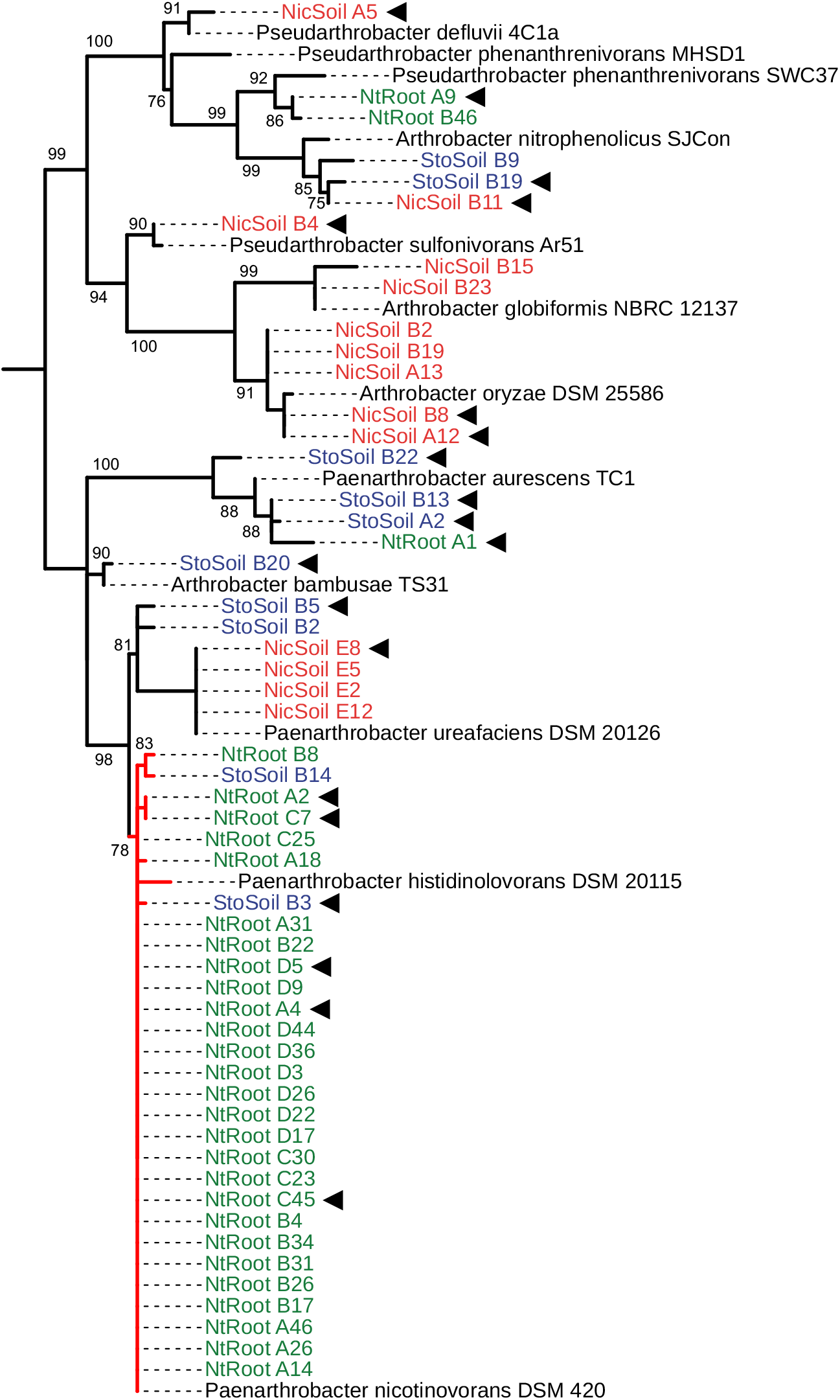
MLE phylogeny of *Arthrobacter* isolates based on their near-complete *16S* rRNA gene sequences. The phylogenetic tree was constructed by an MLE method. Bootstrap values (1000 replicates) above 0.6 are shown in nodes. The red branch represents a monophyletic clade containing almost all isolates from tobacco roots as well as the *A. nicotinovorans* DSM420 strain. Text colors represent the origin of bacterial isolates (green, tobacco roots; blue, santhopine-treated soil; red, nicotine-treated soil; gray, retrieved from the public database). The isolates subjected to whole genome sequencing are marked with arrowheads.

**Fig. S5.**
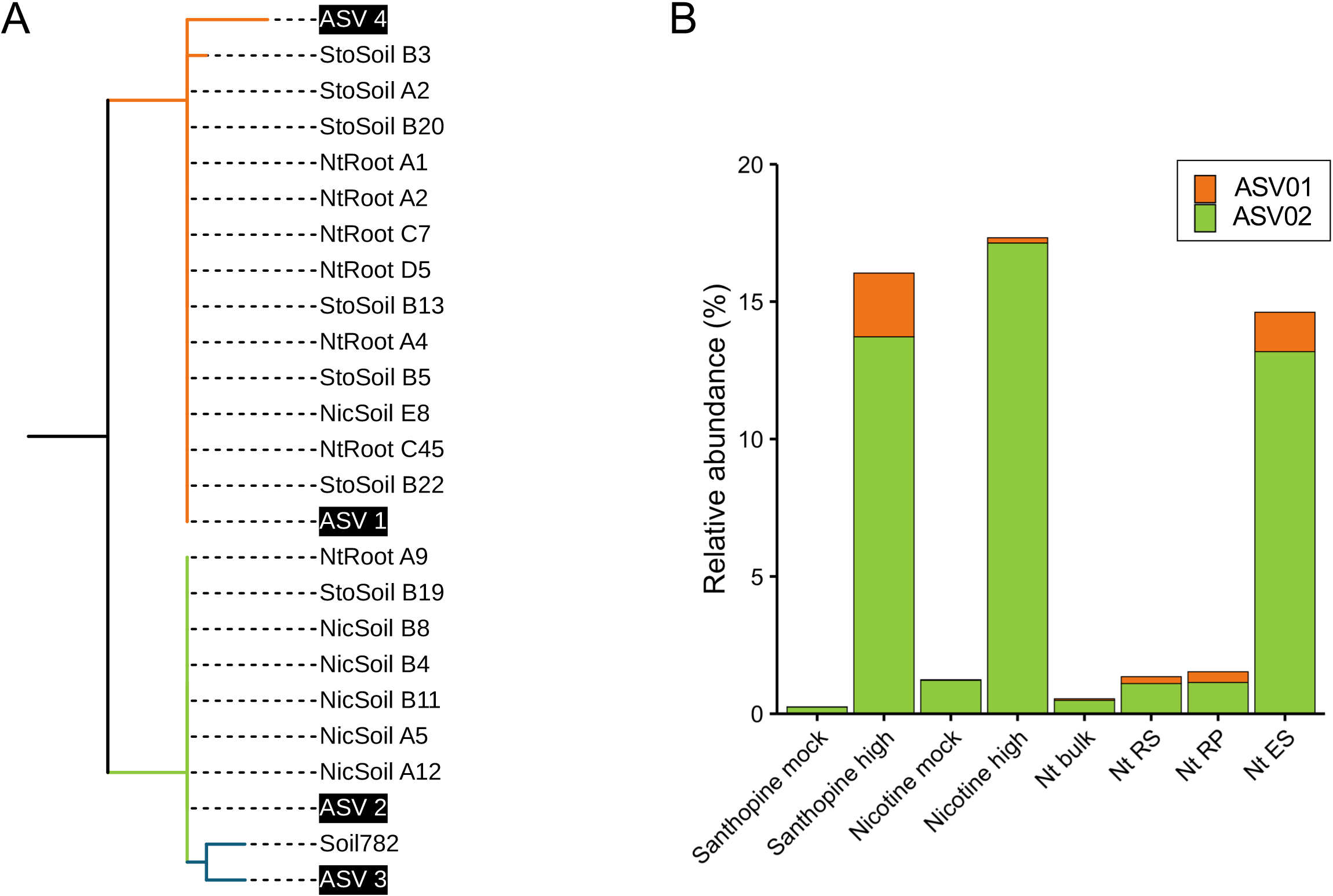
Physiological differences between the sublineages of *Arthrobacter*. **(A)** Representative sequences of the ASVs belonging to the family Micrococcaceae were retrieved and used to infer phylogenetic relationships with the strains isolated from roots and soils using an MLE method. The branch colors represent the sublineages defined by their AMPHORA phylogeny. **(B)** Average relative abundance of ASVs in each sample. Each bar represents the mean relative abundance (n = 3 for tobacco roots and santhopine-treated soil and n = 4 for nicotine-treated soil).

**Fig. S6.**
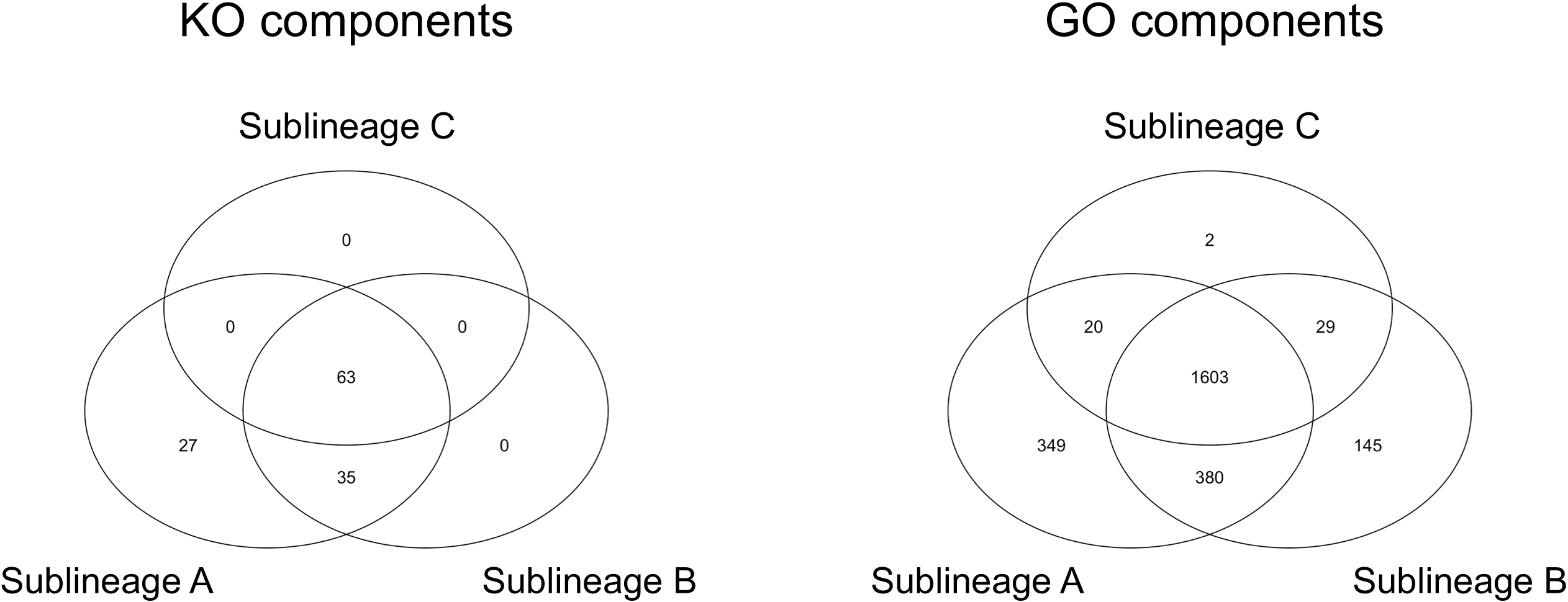
Comparison of ancestral characters of each *Arthrobacter* sublineage. Venn diagrams comparing identified ancestral components based on **(A)** KO and **(B)** OG profiles.

**Fig. S7.**
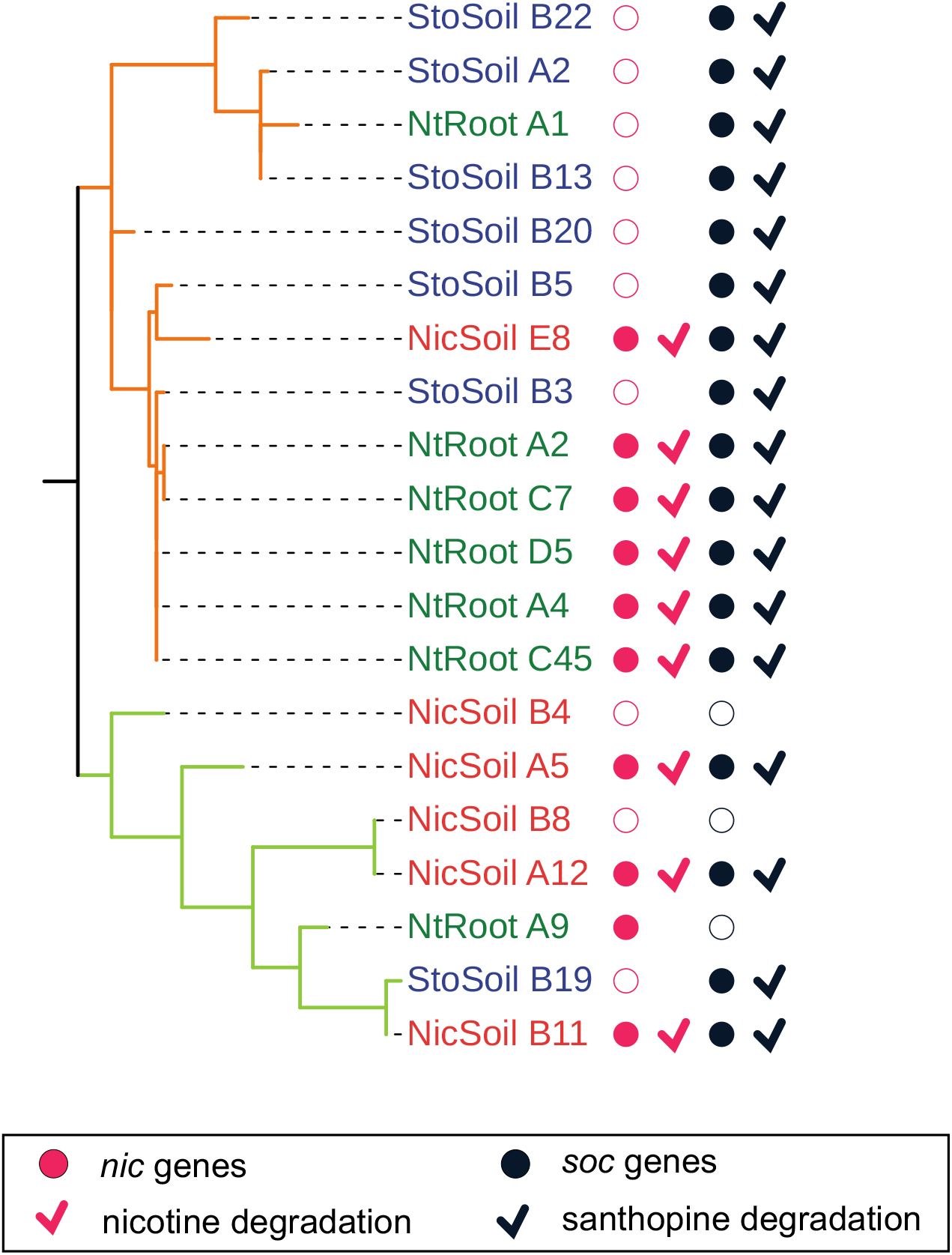
Phylogenetic distribution of catabolic genes and degradation ability within newly sequenced *Arthrobacter*. The presence of catabolic genes and degradation ability of santhopine and nicotine. Text colors indicate their origins. The branch colors represent the sublineages defined by their AMPHORA phylogeny.

**Fig. S8.**
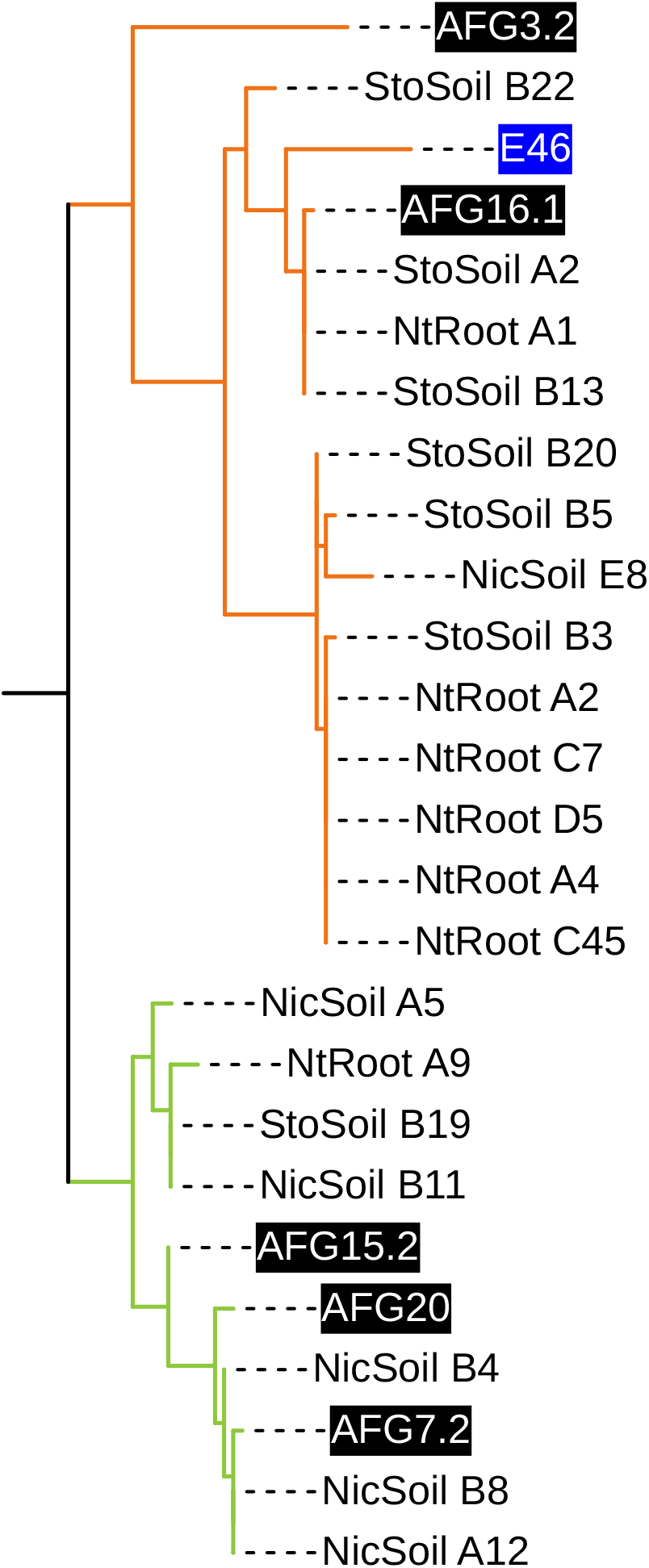
Phylogenetic analysis of plant-derived *Arthrobacter*. *16S* rRNA sequences of AFG20 (KT314111), AFG16.1 (KT314130), AFG7.2 (KT314127), AFG15.2 (KT314122), AFG3.2 (KT314126), and E46 (KT314127) were retrieved from the NCBI. The accession number of each sequence is shown in parenthesis. The MLE method was used to construct a phylogenetic tree with 1000 bootstrap replications. The branch colors represent the sublineages defined by their AMPHORA phylogeny.

**Fig. S9.**
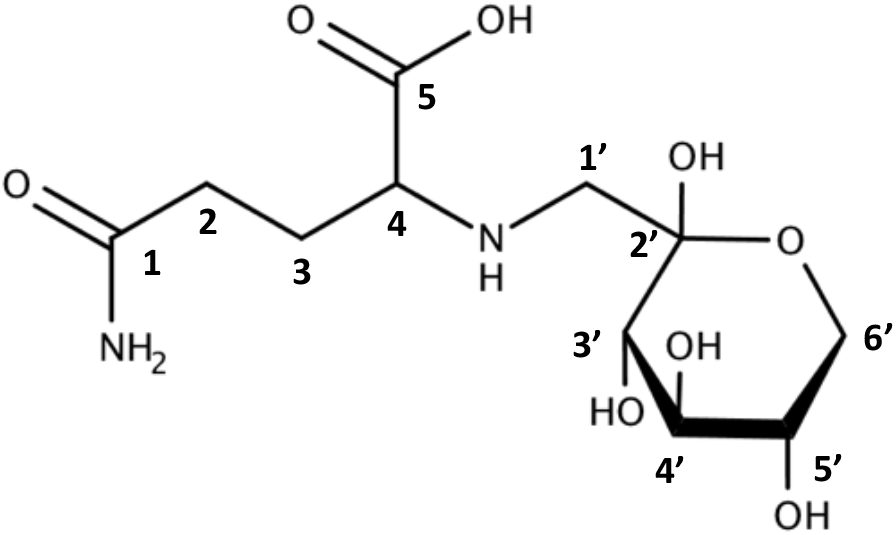
Santhopine structure for NMR analysis.

**Table S1.** NMR data for santhopine.

| Position | Compound A<sup>a</sup> |  | Santhopine<sup>a, b</sup> |  |
| --- | --- | --- | --- | --- |
| | $\delta_C$ (ppm) | $\delta_H$ multiplicity | $\delta_C$ (ppm) | $\delta_H$ multiplicity |
| 1 | 177.8 |  | 177.8 |  |
| 2 | 31.1 | 2.40-2.38 m, 2H | 31.0 | 2.50-2.48 m, 2H |
| 3 | 24.9 | 2.01-1.99 m, 2H | 24.8 | 2.20-2.13 m, 2H |
| 4 | 62.6 | 3.66-3.59 m, 1H (in 3H) | 62.6 | 3.77-3.71 m, 1H (in 3H) |
| 5 | 172.8 |  | 172.7 |  |
| 1' | 52.6 | 3.20-3.18 m, 2H | 52.5 | 3.32-3.25 m, 2H |
| 2' | 95.3 |  | 95.3 |  |
| 3' | 68.9 | 3.92-3.89 m, 1H (in 2H) | 68.8 | 4.02-3.98 m, 1H (in 2H) |
| 4' | 69.9 | 3.76 dd, 1H | 69.8 | 3.77-3.71 m, 1H (in 3H) |
| 5' | 69.3 | 3.66-3.59 m, 1H (in 3H) | 69.3 | 3.85 m, 1H |
| 6' | 63.9 | 3.66-3.59 m, 1H (in 3H) | 63.8 | 3.77-3.71 m, 1H (in 3H) |
|  |  | 3.92-3.89 m, 1H (in 2H) |  | 4.02-3.98 m, 1H (in 2H) |

<sup>a</sup> NMR data for compound A or santhopine in D<sub>2</sub>O
<sup>b</sup> Chen et al. 2016 (16)

**Table S2.** Protein queries for BLASTP search of nic and soc genes.

| Gene | Definition | Accession |
| --- | --- | --- |
| *ndhL* | nicotine dehydrogenase (large subunit) | CAD47952.1 |
| *ndhM* | nicotine dehydrogenase (medium subunit) | CAD47954.1 |
| *6HlnO* | 6-hydroxy-L-nicotine oxidase | CAA11306.1 |
| *Ponh* | 2,6-dihydroxy-pseudo-oxynicotine hydrolase | CAD47941.1 |
| *kdhL* | ketone dehydrogenase (large subunit) | CAD47940.1 |
| *kdhM* | ketone dehydrogenase (medium subunit) | CAD47951.1 |
| *dhph* | 2,6-dihydroxypyridine 3-monooxygenase | CAD47937.1 |
| *socD* | Fructosyl-amino acid oxidase | ABM07460.1 |

**Data Files S1.** Summary data for bacterial isolates from **(A)** tobacco roots, **(B)** santhopine-treated soil, and **(C)** nicotine-treated soil. **(D)**. Composition of bacterial growth media used for isolation approaches.

**Data Files S2.** *16S* rRNA sequences for 54 *Arthrobacter* isolates.

**Data Files S3.** General genomic features of newly sequenced *Arthrobacter* isolates. Assembly qualities were calculated using DDBJ Fast Annotation and Submission Tool (DFAST; https://dfast.nig.ac.jp).

**Data Files S4.** Classification of the isolation sources of *Arthrobacter* genomes for comparative genomics.

**Data Files S5.** Rhizobia genes identified in the genus *Arthrobacter* retrieved from **(A)** *E. meliloti* 1021 (*35*) and **(B)** *R. leguminosarum* bv. *viciae* Rlv3841 (*36*).

**Data Files S6.** List of ancestral components. Ancestral components were estimated based on the KO profile unique to **(A)** sublineage A and **(E)** sublineage A/B. Ancestral components were estimated based on the OG profile unique to **(B)** sublineage A, **(C)** sublineage B, **(D)** sublineage C, and **(F)** sublineage A/B.

